# Altered Cohesin Dynamics During Cellular Differentiation

**DOI:** 10.64898/2026.01.01.697310

**Authors:** Magdalena Jawor, Karol Tchorz, Marcin Ostoja-Helczynski, Susannah Rankin, Jacob Kirkland

**Author notes:** absolutely contributed equally (Magdalena Jawor, Karol Tchorz). International Institute of Molecular and Cell Biology, 4 Ks. Trojdena Street 02-109 Warsaw, Poland. Friedrich Miescher Laboratory of the Max Planck Society, Max-Planck-Ring 9, Tübingen 72076, Germany (Karol Tchorz).

## Abstract

The cohesin complex plays essential roles in chromosome organization and gene regulation, yet how cohesin dynamics are controlled during cell-state transitions remains poorly understood. Here, we examined how cohesin regulation is remodeled during the differentiation of mouse embryonic stem cells (mESCs) into the cardiomyocyte lineage using an *in vitro* differentiation system. We found that core cohesin subunits remain broadly stable at the protein level. In contrast, the levels of cohesin regulators, including the cohesin removal protein WAPL and the cohesin stabilizing protein ESCO1, decline sharply despite modest transcript-level changes. The cohesion maintenance factor Sororin was also reduced. To better understand the net effect of these changes on cohesin dynamics, we use live-cell FRAP of RAD21, which revealed increased cohesin mobility in differentiated cells without a change in recovery kinetics, consistent with reduced stable chromatin engagement or redistribution into a chromatin-unbound nuclear pool. To test functional consequences, we generated homozygous degron alleles for *Wapl* and *Esco1* and induced acute degradation using a dTAG-based system. Loss of WAPL altered cell-cycle dynamics in stem cells and produced a characteristic “vermicelli” chromosome phenotype, consistent with abnormally high and lethal cohesin retention on chromatin. Surprisingly, depletion of ESCO1 had no clear impact on viability and cell cycle progression. Notably, despite loss of detectable WAPL protein in the differentiated cell population, we find that WAPL remains functionally required to maintain a viable interphase chromosome organization. Together, these findings identify cohesin regulators, rather than cohesin abundance, as central drivers of changes in cohesin dynamics during differentiation. They further show that even very low levels of WAPL continue to provide critical structural plasticity of chromosomes following cell cycle exit and lineage commitment.

## INTRODUCTION

A central paradox in biology is that developmental programs require changes in gene expression, and yet genomes are identical within every cell. Much effort has been exerted to understand how changes in gene expression can be accomplished through epigenetic modulation, while maintaining a level of plasticity and responsiveness. The cohesin complex, through its capacity to fold and stabilize the genome into chromosome loops and domains, plays critical roles in gene regulation throughout development (Bonev and Cavalli, 2016; De Koninck et al., 2020; Dowen et al., 2013; Solé-Ferran and Losada, 2025). The topology of the genome imposed by cohesin ensures proper apposition of genes and their regulatory elements, thereby enforcing cell identity (Bryan et al., 2025; Davidson et al., 2019; Dowen et al., 2014; Guacci et al., 2019; Seitan et al., 2013), although other models have been proposed (Hsieh et al., 2022). The indispensability of cohesin in gene expression and development is illustrated by the spectrum of cohesinopathies - multisystem developmental disorders caused by mutations in cohesin subunits and their regulators (Garcia et al., 2021; Mfarej et al., 2023; Solé-Ferran and Losada, 2025; Watrin et al., 2016; Zakari et al., 2015). How, then, is genome topology modified to allow changes in gene expression in response to emerging or changing developmental programs?

Expression of the cohesin complex is essential and occurs in both dividing and post-mitotic cells. Cohesin both forms and stabilizes chromosome loops and domains, and tethers the products of DNA replication together in actively dividing cells in mechanistically distinct pathways (Bender et al., 2020; Lafont et al., 2010; Sansam et al., 2018; Srinivasan et al., 2020). The interaction of cohesin with chromatin is controlled by a number of proteins including variant subunits that modify the loading, stable binding, and removal of the complex from chromosomes (Cuadrado et al., 2022). These include the NIPBL/MAU2 complex that loads cohesin on chromatin, the WAPL protein that removes cohesin from chromatin, and the ESCO1 and ESCO2 acetyltransferases that render cohesin WAPL-resistant (Rittenhouse and Dowen, 2024). Other proteins include the cohesion-maintenance protein Sororin, which stabilizes cohesion between sister chromatids (Rankin et al., 2005), and the HDAC8 deacetylase, which removes ESCO-dependent acetylations, thereby enabling cohesin recycling (Deardorff et al., 2012). Importantly, the interaction of cohesin with chromatin is largely dynamic. A small fraction of cohesin (∼20%) is stabilized by activities associated with the replication machinery to tether sister chromatids together in G2 in proliferating cells (Gerlich et al., 2006). The remaining cohesin associates dynamically with chromatin throughout interphase, with residence times of 15-25 minutes for interphase cohesin to several hours for replication-coupled cohesin (Gerlich et al., 2006; Hansen et al., 2017; Kueng et al., 2006). The net result of this constant cohesin binding, loop formation, and removal is the generation of stereotypical chromosome topology, which can be most readily revealed by chromosome conformation capture techniques (Meluzzi and Arya, 2020; Rao et al., 2017). How then is cohesin binding and stability modulated in response to developmental signals that cause transcriptional changes and altered cell identity? Does the stability of cohesin binding impose cell-type-specific gene regulation? If so, does it change only locally near relevant genes or throughout the genome? To begin to address these questions, we have investigated the changes in cohesin regulation during *in vitro* cell differentiation.

Embryonic stem cells can be induced to differentiate into different cell types *in vitro* by altering the culture substrate and media. Here, we induced a mouse embryonic cell line to transition from pluripotency into the mesodermal lineage, ultimately forming contracting cardiomyocytes, and assessed cohesin regulation during this transition. We found that the level of the core cohesin complex was largely unchanged over the course of the experiment. In contrast, we found a strong decrease in the levels of certain cohesin regulators. Strikingly, we found that the dynamics of cohesin binding changed dramatically following the differentiation protocol. We hypothesize that transcriptional changes that promote cell differentiation may be induced, at least in part, by alterations in cohesin dynamics that affect chromosome structure. These changes in cohesin’s association with chromatin might alter the stability of interactions between genes and their regulatory elements, thereby promoting lineage commitment and stabilizing changes in cell identity.

## RESULTS

### *In vitro* differentiation: the cardiac lineage

Cohesin controls genome folding and is thought to have broad impacts on gene expression and thus cell fate. This, in turn, implies that cohesin is likely to be remodeled in specific ways during cellular differentiation. To define how cohesin is regulated during cellular differentiation, we used an *in vitro* model in which mouse embryonic stem cells (mESCs) differentiate into the mesodermal lineage following a specific series of physical and media manipulations [Fig. 1A] (Lynch et al., 2018). In this method, undifferentiated mESCs are collected as a single-cell suspension and hung in drops from the lids of cell culture plates for 2 days, resulting in the formation of embryoid bodies (EBs). The EBs are then transferred to gelatin-coated plates in differentiation media and allowed to grow out and further differentiate. Under this protocol, cells progress through mesodermally committed stages, then into cardiac precursors, and ultimately into cardiomyocytes.

**Figure 1.**
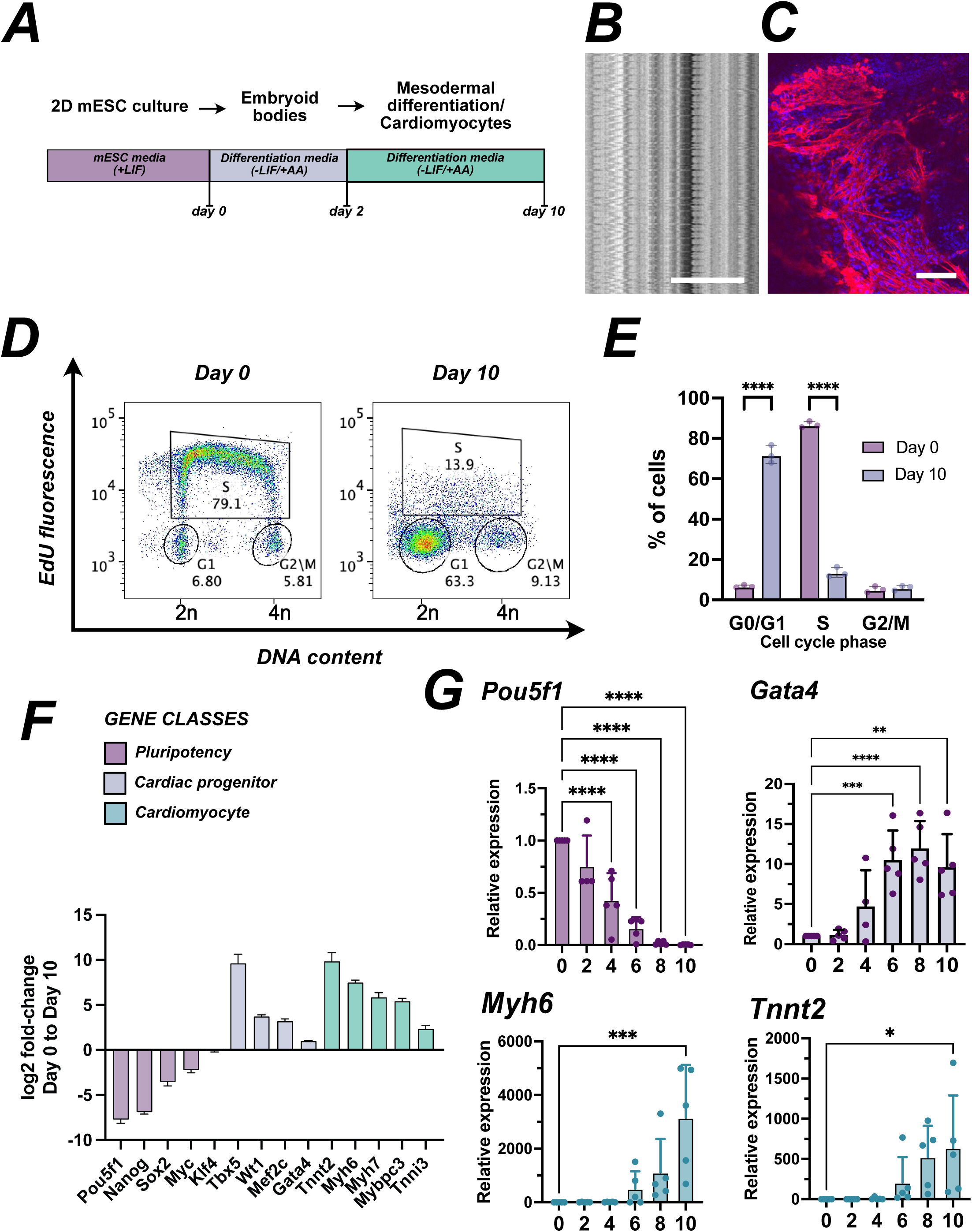
Gene regulation during cell differentiation. **A.** A schematic illustrating the differentiation protocol used in this study. Mouse embryonic stem cells grown in stem cell media containing leukemia inhibitory factor (LIF) were suspended in hanging drops for 2 days in differentiation media (no LIF, supplemented with ascorbic acid). Cells were then plated in differentiation media for up to 8 additional days. indicated. **B. Cells undergo periodic contraction.** Shown is a representative kymograph made from a time-lapse phase-contrast movie after 8 days of differentiation, in which time is on the y-axis. Each contraction resulted horizontal striations over time. **C. Differentiation protocol leads to the formation of cardiomyocytes.** Cells cultured as in **A** were fixed and immunostained with anti-cardiac troponin antibodies and counterstained with DAPI. Patches of elongated cells expressing cardiac troponin were found in all samples. Scale bar = 100μm **D. Flow cytometric analysis of DNA synthesis before and after differentiation.** Mouse ESCs were pulse labeled with EdU before or after the differentiation protocol shown in A, and processed for flow cytometry. Total DNA content is shown on the x-axis; EdU signal is on the y-axis. Gates were drawn as shown to measure G1, S phase, and G2 or M cells. **E. Cell cycle analysis.** Percentages of cells within each gate in panel **D** are plotted. **** = P<0.0001, 2-way ANOVA. **F. Gene expression changes during differentiation.** Bulk RNA from cells isolated either at day 0, or at day 10, were sequenced and analyzed for changes in gene expression of markers relevant to stem cell maintenance and the cardiac lineage. n = 4, error bars show lfcSE. **G. Changes in expression of stem cell markers during differentiation process.** Total RNA samples were collected on the indicated days during the differentiation protocol and analyzed by qPCR for the presence of the indicated mRNAs. Expression levels for all samples were normalized to Llgl1. * P < 0.0332; ** < 0.0021; *** < 0.0002; **** < 0.0001 by Ordinary one-way ANOVA with Dunnett’s multiple comparison test.

We performed several analyses to confirm differentiation in our hands. First, we confirmed that the cells underwent periodic contractions, as seen previously (Lynch et al., 2018). Indeed, by day 8, all cultures showed regions undergoing visible periodic contractions. To document the contractions and measure the frequency, we collected representative time-lapse phase-contrast image series at low magnification, and generated a kymograph [Fig. 1B]. In the example shown, the contraction frequency was ∼ 51.6/sec.

The cardiac troponin T gene (*Tnnt2*) is uniquely expressed by cardiomyocytes, and the encoded protein is required for cardiac muscle contraction (Sehnert et al., 2002). We immunostained our differentiated cells to assess cardiac troponin T protein expression [Fig. 1C], and found that expression of cTnT protein, though very consistent (occurring in ∼100% of individual cultures), did not occur in all cells. In all cultures tested, large patches of cTnT-positive elongated cells were observed, as previously reported (Lynch et al., 2018) [Fig. 1C].

Differentiation typically results in exit from the cell cycle and reduced cell division (Liu et al., 2019). To define the cell cycle distribution of our differentiated cells, we performed 2-dimensional flow cytometry using propidium iodide to measure total DNA content, combined with pulse-labeling with the thymidine analog EdU (5-ethynyl-2’-deoxyuridine) to assess active DNA replication [representative sample Fig. 1D]. Strikingly, prior to differentiation, the majority of the ESCs (86.8%) were in S-phase with 6.7% in G0/G1 and 5.2% in G2/M [Fig. 1E]. In contrast, following the differentiation protocol, cells were primarily in G0/G1 (72.0%), with only 13.6% in S-phase and 6.0% in G2/M. To rule out the possibility that cell death accounts for the differentiation protocol’s effect on cell cycling, we also assessed cell viability in both cultures. We did not observe an increase in cell death at day 10 compared to undifferentiated mESCs [SF1A]. We conclude from this experiment that cells remain viable following the differentiation protocol, which results in exit from the cell cycle consistent with differentiation studies in other models and cell types (Calder et al., 2013; Coronado et al., 2013; Walsh and Perlman, 1997).

Following the differentiation protocol, our results indicate that most cells have exited the cell cycle and that many have differentiated into cardiomyocytes. To better understand changes in cell identity, we performed bulk RNA-seq on mESCs and compared the resulting transcriptome with that of the differentiated cell population on day 10. Genes from different stages of differentiation were selected from (Hota et al., 2022) and (Wamstad et al., 2012). Consistent with differentiation and loss of stemness, we found that several pluripotency factors, including *Pou5f1 (Oct4)*, *Nanog, Sox2*, and *Myc,* were down-regulated in the day 10 sample (fold-change < 0.5 and adj p-value < 0.05), though the expression of the pluripotency marker *Klf4* remained unchanged (log2FC −0.13; fold-change 0.92; adj p-value 0.15) [Fig 1E, SF1B]. Interestingly, mesodermal genes showed both up-and down-regulated transcripts at day 10 [SF1C]. Cardiac progenitor genes, including *Tbx5, Wt1, Mef2c*, and *Gata4*, were all significantly up-regulated (fold-change > 1.5 and adj p-value < 0.05) [Fig 1E, SF1D]. The cardiomyocyte-specific genes *Tnnt2, Myh6, Myh7, Mybpc3*, and *Tnni3* were all up-regulated in differentiated cells compared to mESCs (fold-change > 1.5 and adj p-value < 0.05) [Fig 1E, SF1E]. We conclude from these experiments that the protocol here results in loss of stemness and exit from the cell cycle.

To assess the kinetics of the observed transcriptional changes during differentiation, we isolated RNA at 2-day intervals. Using RT-qPCR, we quantified messages for *Pouf1* (Niwa et al., 2000) (a pluripotency marker), *Gata4* (Heuvelmans et al., 2025) (a transcription factor expressed in cardiac progenitor cells), and *Myh6* and *Tnnt2 (Veevers et al., 2018)* (cardiomyocyte markers) [Fig. 1F]. We found that *Pou5f1* transcripts steadily decreased to near undetectable levels by day 10. In contrast, *Gata4* transcripts increased from day 4 onward (significant at days 6, 8, and 10) but began to decline at day 10 [Fig. 1F]. Consistent with the development of cardiomyocytes, the abundance of *Myh6* and *Tnnt2* transcripts increased (mean value 3159-fold and 640-fold, respectively) throughout the differentiation protocol [Fig. 1F].

### Expression of cohesin regulators during differentiation

Because we are interested in the role of the cohesion apparatus in cell differentiation, we analyzed the expression of both core cohesin subunits and cohesin regulators during *in vitro* differentiation. Comparing bulk RNAseq results from mESCs with differentiated cells (day 10) indicated that transcription of core cohesin subunits *Smc1a* was unchanged (log2FC 0.21; fold-change 1.15; adj p-value 0.07), or modestly reduced *Smc3* (log2FC −0.67; fold-change 0.63; adj p-value 1.26e^−6^) and *Rad21* (log2FC −0.76; fold-change 0.59; adj p-value 2.16e^−21^) [Fig. 2A]. Transcript levels of *Nipbl*, which loads cohesin onto chromatin and modulates chromosome loop extrusion, were also unchanged (log2FC = 0.105; fold-change 1.07; adj p-value = 0.28). Transcripts for the cohesin releasing factor *Wapl* (log2FC −0.93; fold-change 0.52; adj p-value 1.39e^−33^) and the cohesin stabilizing protein *Esco1* (log2FC −0.64; fold-change 0.64; adj p-value 3.18e^−8^) were modestly down-regulated. In contrast, transcripts for the cohesion maintenance proteins *Esco2* (log2FC −2.25; fold-change 0.21; adj p-value 4.36e^−70^) and *Cdca5* (Sororin: log2FC −1.75; fold-change 0.29; adj p-value 5.45e^−48^) were strongly down-regulated [Fig. 2A]. Consistent with exit from the cell cycle, *Cdca5* (sororin) is a direct target of E2F1, a master regulator of cell cycle entry (Chen et al., 2019). Transcripts of *Hdac8*, which deacetylates SMC3, were elevated in differentiated cells compared to stem cells (log2FC 1.20; fold-change 2.30; adj p-value 4.29e^−6^) [Fig. 2A]. Cell cycle-associated genes, *Ccne1* (Cyclin E1; log2FC −3.22; fold-change 0.11; adj p-value = 3.66e^−156^), *Ccnb1* (Cyclin B; log2FC −2.66; fold-change 0.16; adj p-value = 4.20e^−11^), *Ccnd1* (Cyclin D1; log2FC −1.02; fold-change 0.49; adj p-value = 1.73e^−6^), and *Aurkb* (Aurora B; log2FC −0.96; fold-change 0.51; adj p-value = 2.19e^−19^) were all decreased in day 10 differentiated cells compared to mESCs [Fig. 2A].

**Figure 2.**
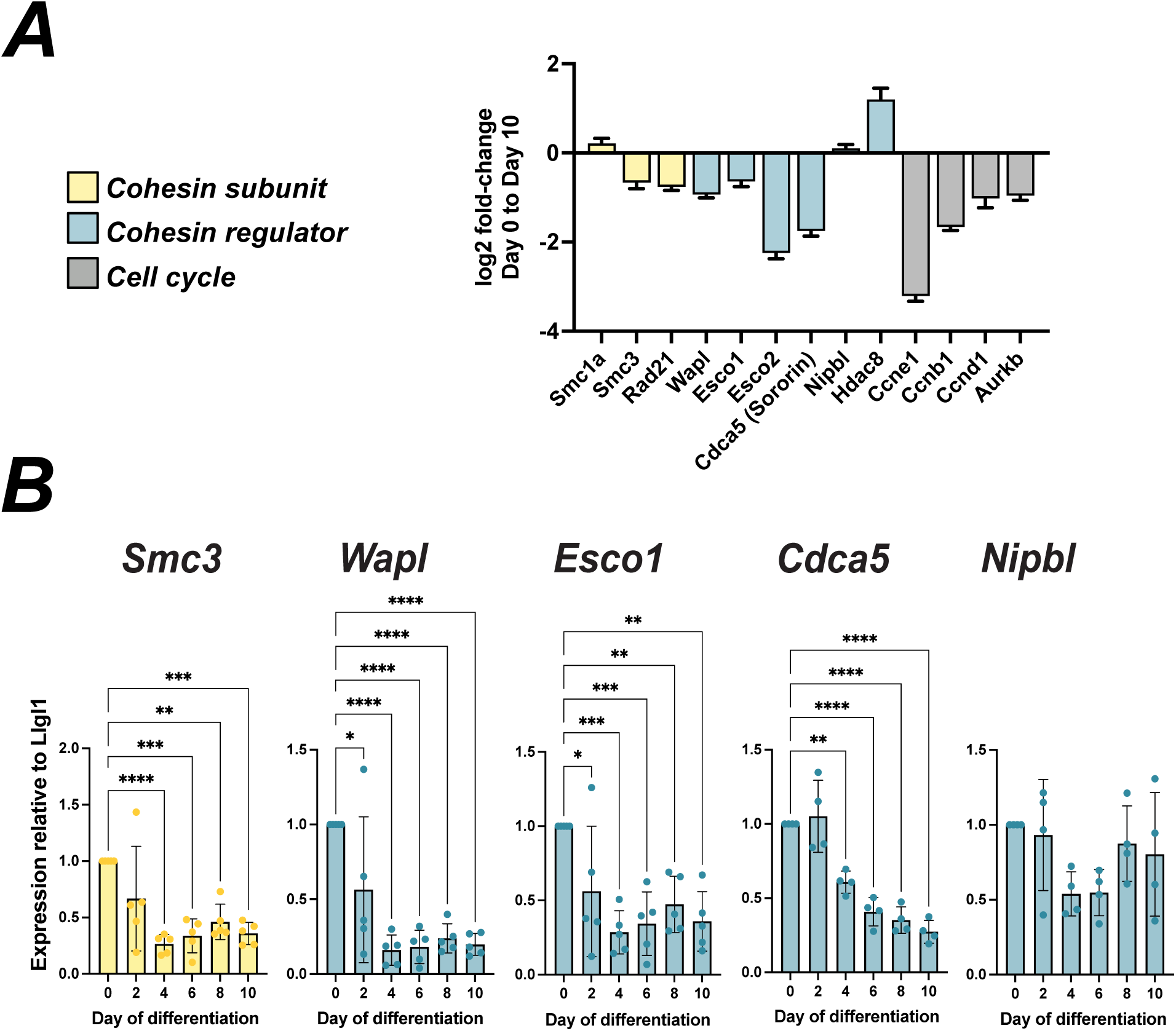
Expression of cohesin regulators during *in vitro* differentiation. **A. Changes in gene expression in differentiated cells.** The relative expression of the indicated cohesin regulators were compared between TC1 cells and day 10 of differentiation. n = 4, error bars show lfcSE **B. RNA levels.** mRNA levels of the indicated genes were assessed by qPCR over the full course of *in vitro* differentiation. All expression levels were analyzed relative to Llgl1, which remained constant. * = p< 0.0332; ** = p< 0.0021; *** = p< 0.0002; **** = p< 0.0001 by Ordinary one-way ANOVA with Dunnett’s multiple comparison test.

Detailed qPCR analysis through the differentiation time course confirmed that transcripts encoding SMC3 decreased by ∼50% beginning at day 4 and persisting at this reduced level through day 10 [Fig. 2B]. *Nipbl* transcripts were not significantly altered at any time during differentiation. In contrast, *Wapl* and *Esco1* transcripts were reduced as early as day 2, maximally downregulated by day 4, and persisted at this reduced level through day 10. *Cdca5* transcripts were down-regulated at day 4, with further decreases through day 10 [Fig. 2B].

Because transcript levels do not always reflect protein levels, we also assessed protein levels by immunoblot analysis. Despite reductions in *Smc3* transcripts detected by both RNAseq and RT-qPCR, SMC3 and RAD21 protein levels remained relatively stable throughout the 10-day differentiation [Fig. 3A-B]. In contrast, the levels of the WAPL and ESCO1 proteins decreased to an even greater extent than their respective transcripts. WAPL protein levels were reduced to undetectable levels at day 8 and day 10 [Fig. 3A-B]. ESCO1 protein levels were significantly reduced as early as day 2 and became nearly undetectable by day 8. Sororin levels were significantly decreased by day 2 and continued to decline further through day 10. Finally, HDAC8 protein levels appeared to increase throughout differentiation, similar to the pattern of *Hdac8* transcripts [Fig. 3A-B]. Because we observed reduced cell growth and division, we also measured the levels of the major mitotic cyclin, Cyclin B1 (*Ccnb1*). Consistent with cells exiting the cell cycle during differentiation, Cyclin B1 protein levels were reduced as early as day 2 and continued dropping through day 10. [Fig. 3A-B].

**Figure 3.**
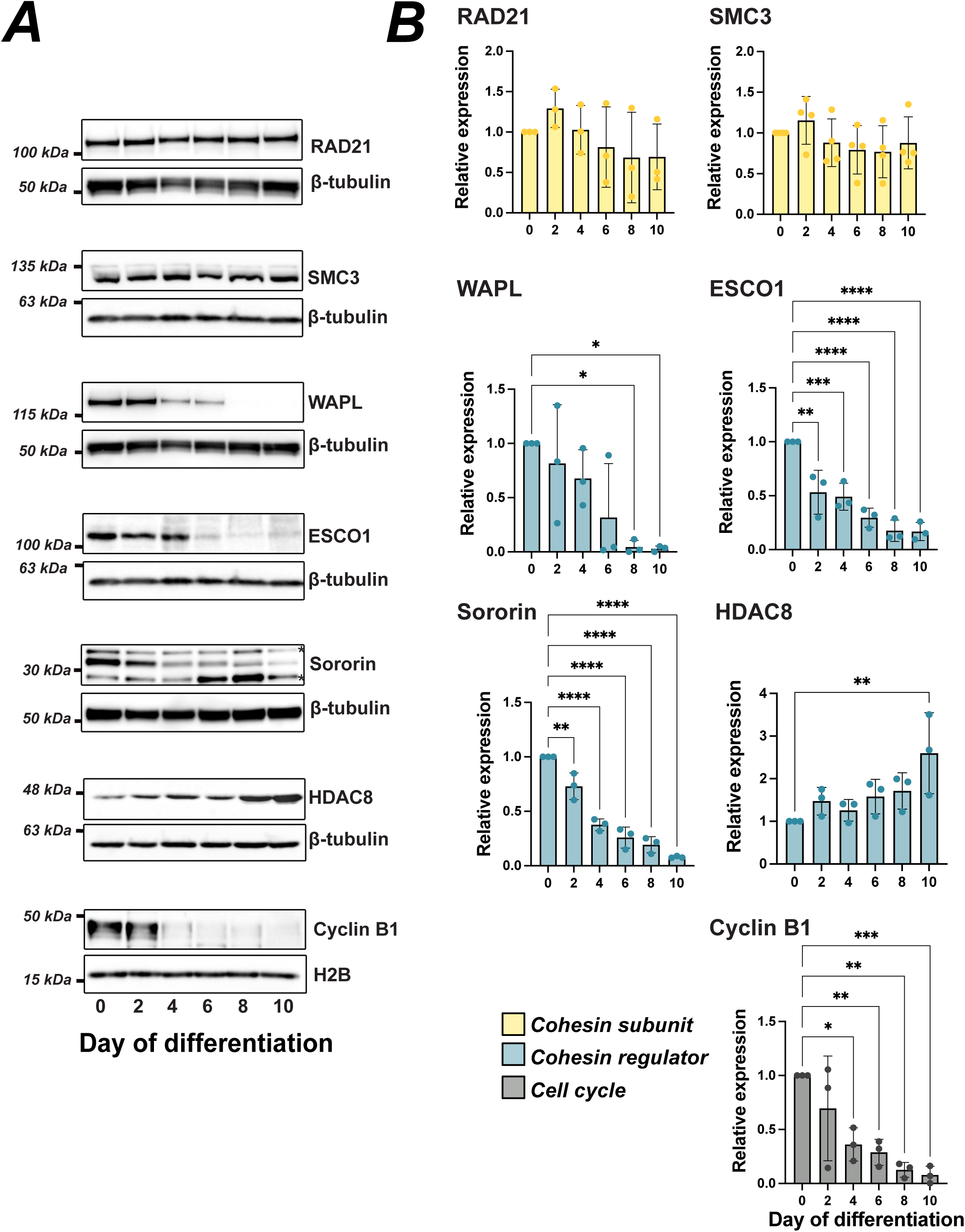
Protein levels of cohesin regulators during *in vitro* differentiation. **A. Immunoblot analysis.** Whole cell lysates collected on different days during the differentiation protocol were resolved by SDS-PAGE and probed with antibodies against the indicated proteins. β-tubulin and histone H2B were used as a loading control. **°** denotes non-specific bands with the Sororin antibody. **B. Aggregate data.** The experiment shown in **A** was repeated for a total of at least three replicates, and protein levels were quantified relative to day 0, using β-tubulin or H2B as a loading control. For each protein n 3, * = p< 0.0332; ** = p< 0.0021; *** = p< 0.0002; **** = p< 0.0001 by Ordinary one-way ANOVA with Dunnett’s multiple comparison test.

### WAPL and ESCO1 degradation in mESCs

The marked decrease in WAPL and ESCO1 protein levels during the differentiation protocol suggested that differentiation may, in part, unfold through changes in the stability of chromatin binding by cohesin. We therefore tested whether loss of either protein, both of which control the stability of cohesin association with chromatin, might impact the differentiation process. To achieve this, we generated derivatives of the TC1 cell line that enable acute and reversible depletion of either WAPL or ESCO1 protein in two separate cell lines. We selected the VHL-based dTAG^V^-1 system for its superior performance in differentiated cells, which is essential for subsequent experiments using the *in vitro* differentiation model (Nabet et al., 2020). The FKBP12^F36V^ tag is functionally inert under basal conditions but induces rapid protein degradation upon treatment of the cells with the dTAG^V^-1 ligand, which promotes interaction of the tagged proteins with the VHL E3 ubiquitin ligase, triggering ubiquitination of the target protein and subsequent clearance by the 26S proteasome. Using CRISPR–Cas9–mediated genome editing to introduce the FKBP12^F36V^ coding sequence, followed by clonal isolation and PCR-based genotyping, we identified homozygous lines [Fig. 4A; SF2A]. We reasoned that this system would allow high temporal resolution in studying the impacts of WAPL and ESCO1 on chromatin behaviors while also minimizing compensatory effects associated with stable genetic perturbation.

**Figure 4.**
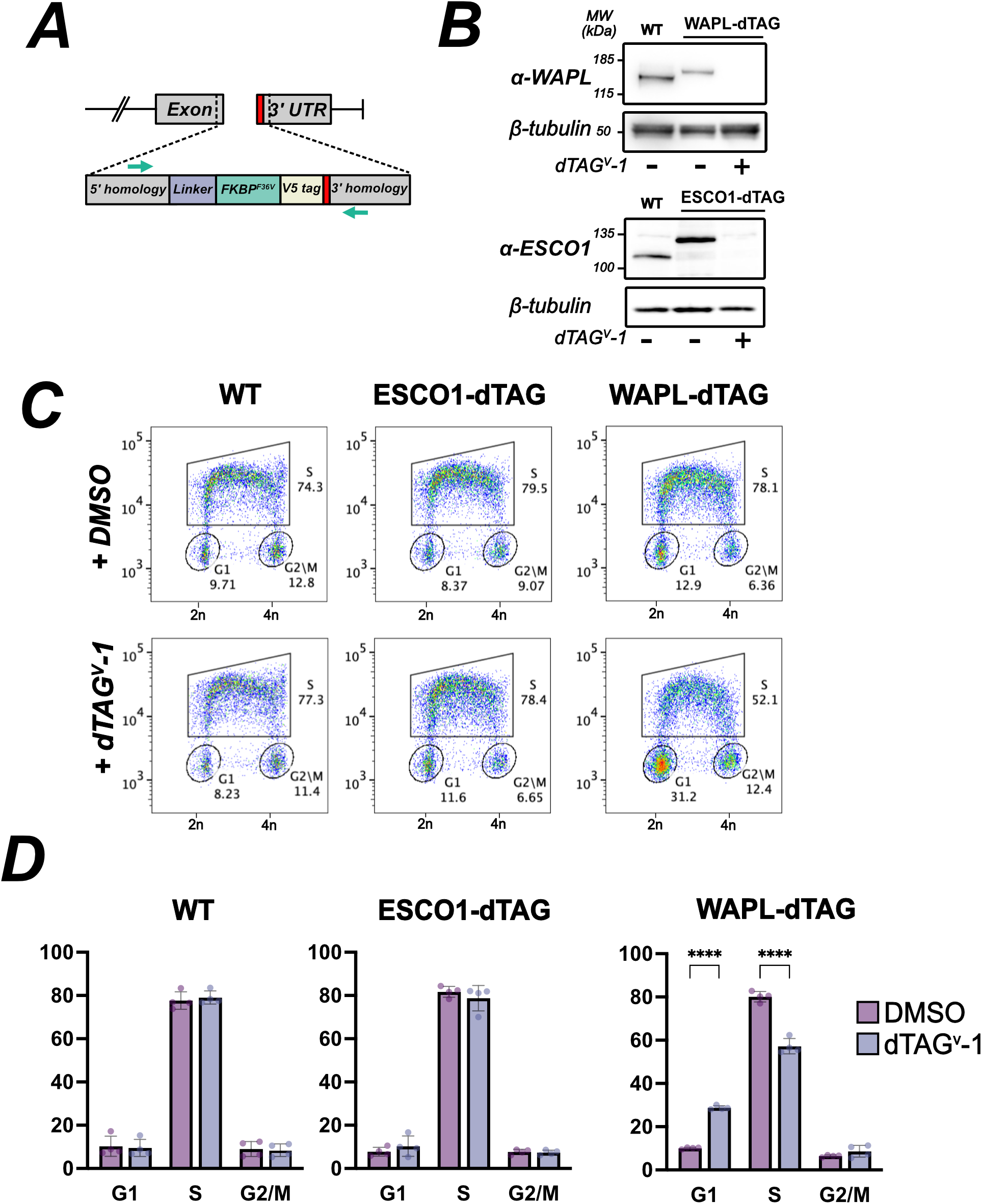
Construction and characterization of degron cell lines. **A. Strategy.** DNA double-strand breaks were induced near the stop codon (red) and a cassette with gene homology flanking the break site (gray) was introduced. Included in the cassette were sequences encoding a flexible linker (lavender), the FKBP^F36V^ allele (green) and the V5 epitope tag. Green arrows indicate position of PCR primers used to validate insertion at endogenous genomic loci (Fig. S2A). **B. Validation of degrons.** Immunoblot of WAPL and ESCO1 in degron cell lines. Cells with homozygous tagged alleles showed reduced protein mobility and loss of signal following 24h treatment with dTag^v^-1. **D. Loss of WAPL causes reduced cell cycling.** Control (TC1) or degron cells (ESCO1-dTag or WAPL-dTag) were treated with dTAG-V1 for 24 (Fig. S2C) or 48 hours (shown) and DNA synthesis was assessed by pulse labeling with EdU, as in Figure 1. Gates were drawn to include cells in G1, S phase, or G2, and these numbers are shown graphed as a percent of the cell population in panel C. **** = p < 0.0001 by 2-way ANOVA.

We confirmed the presence of the dTAG and that levels of ESCO1 and WAPL were sensitive to treatment with the dTAGv-1 ligand. Indeed, the mobility of both ESCO1 and WAPL was reduced by approximately 15 kDa relative to the parental cell line, consistent with the presence of the tag, and we observed no evidence of residual untagged protein, consistent with bi-allelic modification. After 24h treatment with dTAG^V^-1 the levels of each protein were significantly reduced [Fig. 4B]. We noted that the dTagged WAPL was expressed at a lower level than in wild-type cells, even in the absence of dTAG^V^-1 treatment [Fig. 4B]. Conversely, tagged ESCO1 was expressed at a slightly higher level than the untagged protein in ESCO1 in the parental cell line [Fig. 4B]. Both WAPL and ESCO1 were degraded to undetectable levels after 24h treatment with dTAGv-1 [Fig. 4B].

In some cell types, WAPL has been shown to be essential, while other studies using conditional protein degradation suggest that WAPL may be dispensable under some conditions (Liu et al., 2025; Tedeschi et al., 2013). The ESCO1 protein has been shown to affect chromosome-loop homeostasis, but its impact on cell growth and proliferation is less clear (Wutz et al., 2020). To assess the impact of these proteins on cell growth in our system, we performed 2-dimensional flow cytometry as described above. Comparing WT, WAPL-dTAG, and ESCO1-dTAG lines under DMSO (vehicle) treated conditions, we found that their cell cycle profiles were similar, consistent with minimal impact of the tagged alleles on cell cycle progression [SF2B]. In contrast, the WAPL-dTAG cells showed a significant shift in the cell cycle profile after 48 hours of dTAG^V^-1 treatment, but not after 24 hours. These cells showed a relative increase in G0/G1 cells (28.8% versus 10.1% in DMSO controls) and a decrease in S-phase population (57.1% versus 80.1% in DMSO controls) [Fig. 3C-D] [SF2C]. Consistent with this, we detected a growth defect in WAPL-dTAG cells after 4 days of treatment by CellTiter-Blue viability assay, which measures metabolic activity (Promega Corp.) [SF2D]. The observed result is consistent with previous work in mESCs, in which co-depletion of WAPL and CTCF had no effect on proliferation or cell cycle during the first 24 hours but led to detectable changes at later time points (Liu et al., 2025). Degradation of ESCO1-dTAG had no significant effect on cell-cycle profiles over a similar time course. We conclude from these experiments that cell proliferation is greatly inhibited by WAPL degradation. In fact, we were unable to sustain cultures of the WAPL-dTAG cell in the presence of dTAG^V^-1.

Acute loss of WAPL in interphase cells produces a characteristic “vermicelli” chromosome phenotype, marked by detectable thread-like chromosomes and abnormal chromatin compaction (Rhodes et al., 2017; Tedeschi et al., 2013). Because WAPL is required for cohesin release, its loss leads to aberrant retention of cohesin along the length of the chromosomes, resulting in a condensation-like phenotype with enhanced chromosome binding of cohesin, and retention of cohesin on mitotic chromosomes from which it is normally largely released in mitotic prophase (Haarhuis et al., 2013; Waizenegger et al., 2000). These changes likely distort higher-order genome organization, impairing the dynamic cohesin binding that normally occurs in interphase, and perhaps promoting abnormal interchromosomal entanglement. To score the “vermicelli” chromosome phenotype, we immunostained cells depleted for WAPL for the SMC3 subunit of cohesin. Cells exhibited a vermicelli phenotype following WAPL degradation, beginning 24h following treatment with dTAG^V^-1 [Fig. 5A-C]. This phenotype, while most obvious with SMC3 immunostaining, was also evident in DAPI-stained nuclei. Additionally we find that SMC3 remains on mitotic condensed chromosomes following WAPL degradation in contrast to untreated cells (Haarhuis et al., 2014; Samejima et al., 2024) [Fig. 5B]. We confirm with these observations that WAPL-dependent cohesin release is essential to maintain proper chromosome architecture, and that depletion of the protein using the dTAG system is sufficient to generate the vermicelli phenotype. This phenotype of altered chromosome structure is likely the proximal cause of poor growth and viability in these cells, as seen previously (Tedeschi et al., 2013).

**Figure 5.**
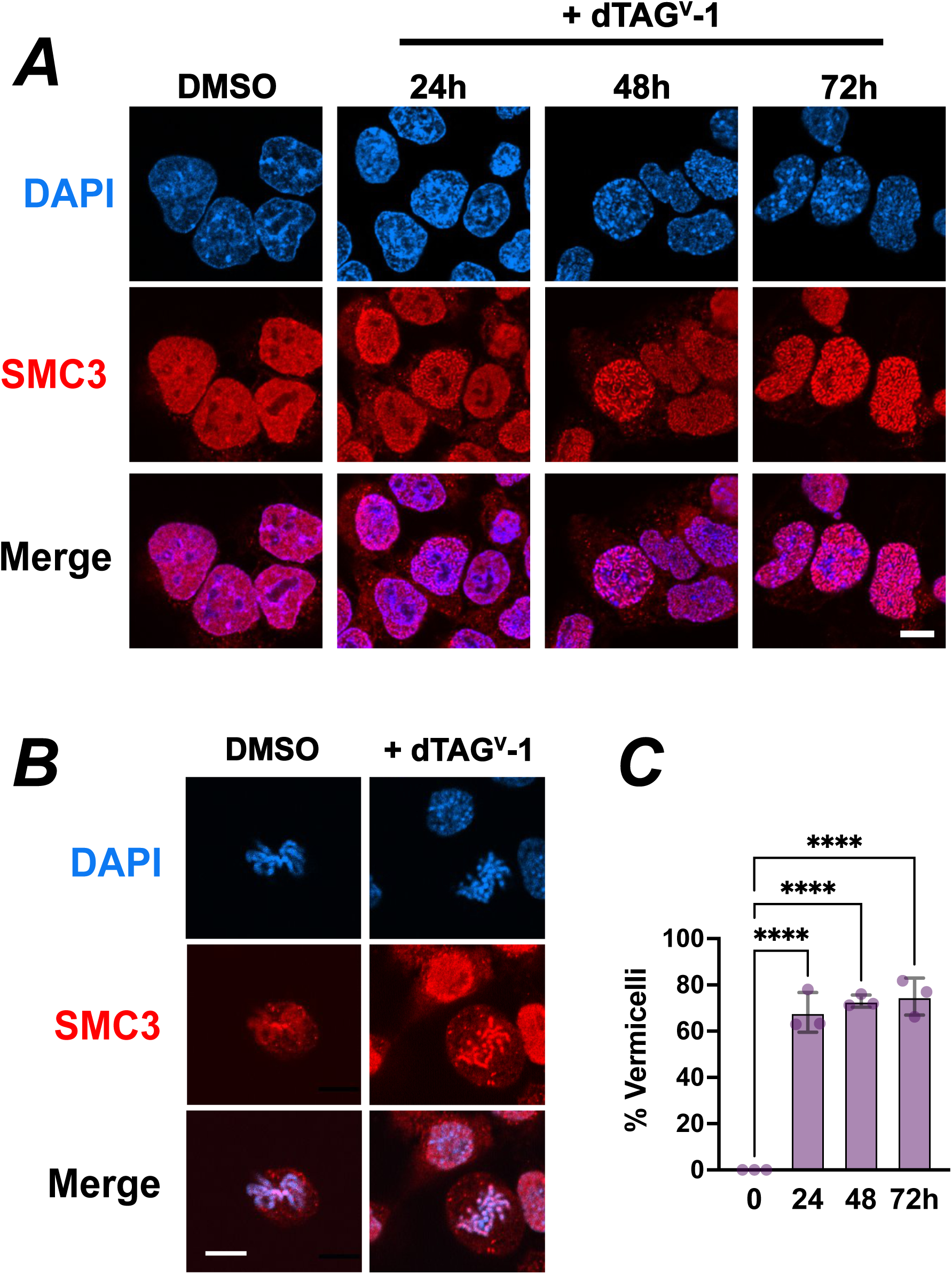
Depletion of WAPL leads to the vermicelli phenotype in stem cells. **A. Vermicelli chromosomes.** The WAPL-dTAG cell line was treated with dTAG^V^-1 for the indicated times, then fixed and stained with anti-SMC3 antibody. The sham-treated cells were incubated with vehicle (DMSO) for 72 hours. **B. Cohesin retention on mitotic chromosomes.** In control cells, cohesin is largely removed during mitotic chromosome condensation (left). Following WAPL depletion cohesin is retained om mitotic chromosomes (right). **C. Quantification of phenotype.** The percent of nuclei showing the vermicelli phenotype, that is condensed interphase chromosomes based on cohesin staining was calculated for each treatment and the average of three experiments is shown > 100 cells/sample. **** = p < 0.0001 by Ordinary one-way ANOVA with Dunnett’s multiple comparison test.

### Cohesin dynamics change during differentiation as the levels of cohesin regulators change

Because the expression of both positive (ESCO1) and negative (WAPL) regulators of cohesin stability was greatly downregulated during our differentiation protocol, we could not draw conclusions about the net effect of these changes on cohesin dynamics in the differentiated cells. We therefore developed a strategy to assess cohesin dynamics using live-cell imaging of the cohesin core subunit RAD21. We used CRISPR–Cas9–mediated genome editing to introduce the HaloTag coding sequence into both alleles of the *Rad21* gene, followed by clonal isolation from single cells and PCR-based genotyping to identify correctly edited homozygous lines. A puromycin selection cassette included in the donor construct was subsequently excised as previously described (Nora et al., 2020). HaloTag expression was confirmed by immunoblotting, which revealed the expected electrophoretic mobility shift of RAD21 in comparison to the wild-type parental cell line [Fig. 6A]. To assess how differentiation affected protein dynamics on chromatin, we performed fluorescence recovery after photobleaching (FRAP) in live cells. A brief, laser pulse was used to selectively photobleach a defined nuclear region containing HaloTag–RAD21 (approximately half the nucleus), and fluorescence recovery was monitored for 26 minutes by confocal imaging, with the unbleached side of the nucleus acting as a reference [Fig. 6B]. In this technique, recovery kinetics reflect molecular exchange from unbleached nuclear regions and provide a quantitative measure of mobility of the labeled RAD21 protein. While the fluorescence recovery rates were very similar in mESCs and differentiated cells (t½ = 3.256 min vs 3.38 min), we observed that the extent of fluorescence recovery was significantly greater in differentiated cells than in parental ESCs [Fig. 6D]. This suggests that cohesin is generally more mobile in our differentiated cells, that a smaller fraction of cohesin is stably bound, or both.

**Figure 6.**
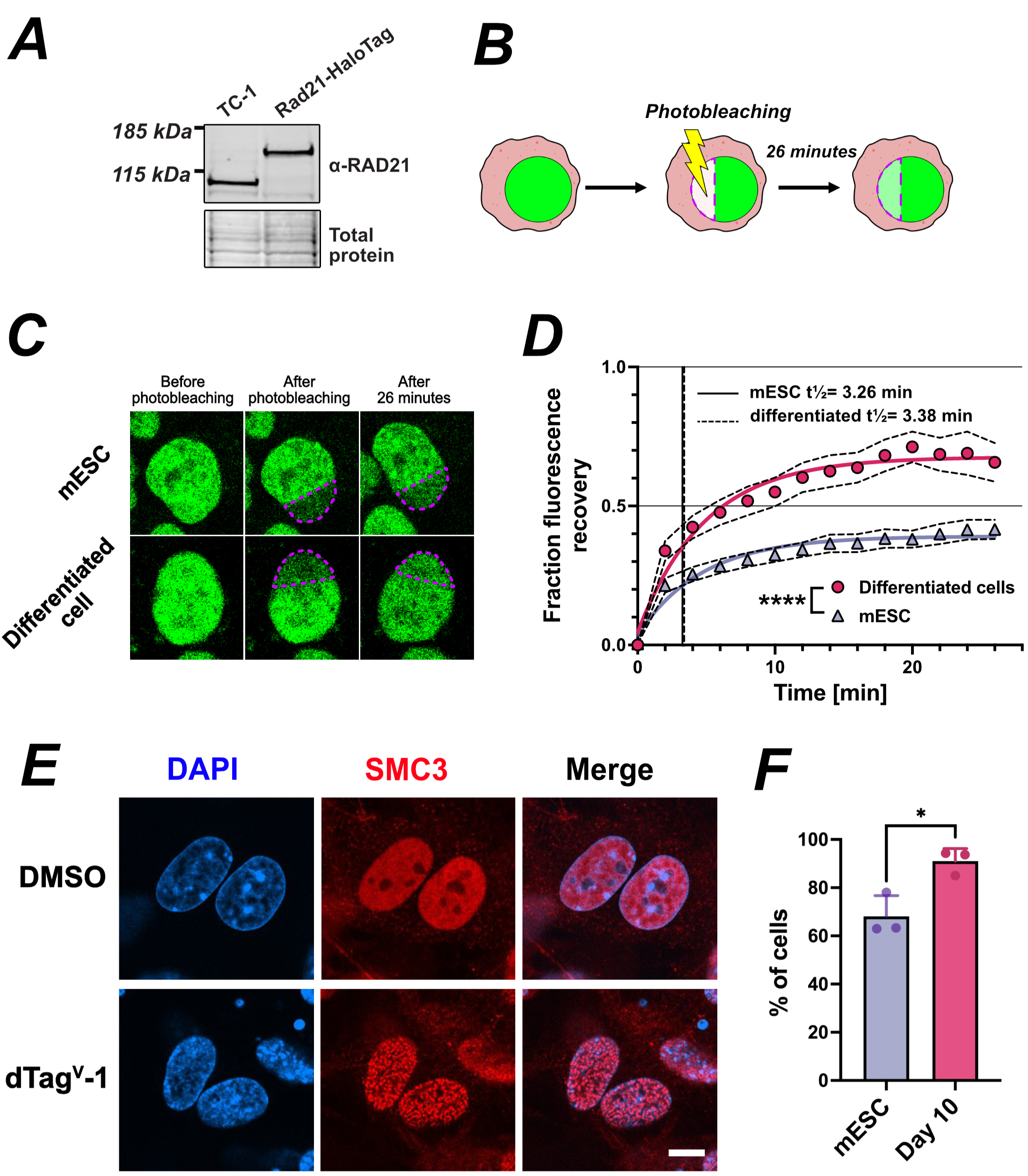
Cohesin dynamics change during differentiation. **A. Halo-tagging of Rad21.** Shown is an immunoblot probed with anti-Rad21 antibody. The parental line, TC1, is shown at left, and the homozygous tagged cell line is shown at right. **B. Fluorescence recovery after photobleaching.** Cells expressing Halo-tagged Rad21 were plated, labeled with Halo-green, and labeled nuclei were irradiated to photobleach signal in ∼1/3 of the nuclear area. Signal recovery in the irradiated part of the nucleus was then scored and normalized to a reference area in the non-irradiated part of the nucleus. **C. Representative images.** Shown are examples of the TC1-Rad21^Halo^ line before and after differentiation for 10 days. Nuclear regions outlined in purple were irradiated and then scored for fluorescence recovery. **D. FRAP analysis of Rad21 dynamics in TC1 and differentiated cells.** The percent fluorescence recovery, normalized to a non-bleached nuclear reference area, is plotted over 26 minutes post-irradiation. The curves represent the average fluorescence recovery following irradiation of n ≥35 nuclei. In stem cells, the recovery plateau reached 39% (fraction 0.39), whereas in differentiated cells, it reached 68% (fraction 0.68). Vertical lines represent the half-time of recovery. Dotted curves represent 95% confidence intervals for each set of measurements, while the solid curves represent the least squares fit of one-phase association. A linear mixed-effects model (REML) showed that fluorescence recovery was significantly affected by time, cell type, and the interaction between the two (**** = p< 0.0001). **E. Differentiated cells form vermicelli upon WAPL depletion.** Cells that were induced to undergo differentiation for 9 days were treated with dTag^V^-1 for 24 hours, fixed, and immunostained with anti-SMC3 antibody as above. **F. Quantification of vermicelli phenotype.** The percent of nuclei showing the vermicelli phenotype was scored after 24 hours of dTag^V^-1 treatment. n = 3 replicates of > 100 cells/sample, * = p <0.05 by unpaired t-test.

Although the level of WAPL protein was below detectable limits following the differentiation protocol, we wondered whether even these low levels of protein continued to provide critical impacts on cohesin dynamics. To address this question, we tested whether further depletion of WAPL in differentiated cells (on day 8 of the protocol) would impact cohesin dynamics. Strikingly, the addition of dTAG^V^-1 was sufficient to induce the “vermicelli” chromosome phenotype, even though the levels of WAPL protein were already greatly reduced [Fig. 6E-F; Fig. 3]. These findings indicate that even the low level of WAPL present in differentiated cells contributes significantly to cohesin dynamics and the maintenance of interphase chromatin organization.

## DISCUSSION

Here, we define how the regulation of cohesin, rather than cohesin abundance, is rewired during stem cell differentiation. Although core cohesin subunits remain largely stable, multiple cohesin regulators, including WAPL, ESCO1, Sororin, and HDAC8, undergo marked changes in expression and abundance during lineage commitment. These transitions are accompanied by altered cohesin dynamics, which in turn likely alter chromatin organization, placing regulatory control of cohesin dynamics at the center of differentiation-associated chromatin change. Our findings indicate that cohesin regulators may be the primary drivers of chromatin dynamics during developmental transitions.

Despite modest transcript-level reductions in core subunits, RAD21 and SMC3 protein levels remained relatively constant during differentiation, suggesting active buffering against loss of the cohesin complex itself, and perhaps reduced expression in the absence of active cell division. The cohesin regulatory proteins WAPL, ESCO1, and Sororin were also moderately reduced at the transcript level. However, in contrast to the cohesin core subunits, they were sharply downregulated at the protein level, with WAPL and ESCO1 becoming nearly undetectable. This discrepancy between transcript and protein levels suggests the presence of a post-transcriptional regulatory layer that targets cohesin modulators during lineage commitment.

Consistent with this idea, acute loss of WAPL in interphase cells resulted in a pronounced “vermicelli” chromosome phenotype characterized by noodle-like condensed chromatin with abnormal compaction. This morphology reflects pathological retention of cohesin on chromatin and excessive stabilization of chromatin loops. This phenotype manifests in interphase, consistent with prior observations that WAPL continuously regulates chromosome topology, not solely at mitotic onset, but throughout interphase (Kueng et al., 2006). Our findings indicate that cohesin release is a central driver of interphase genome organization and support models in which loop extrusion must maintain dynamicity to ensure normal nuclear architecture.

Remarkably, although WAPL protein levels were strongly reduced in differentiated cells, residual WAPL remained functionally essential. Acute chemical degradation in day-9 differentiated cells was sufficient to induce the vermicelli chromosome morphology, demonstrating that even low-abundance WAPL plays an active role in maintaining genome structure post differentiation. These data argue against a simple on/off model and instead support a threshold model, in which small amounts of WAPL buffer against pathological compaction of chromatin. Differentiated cells thus may function near a regulatory boundary, as there is limited redundancy for cohesin release pathways.

Analysis of cohesin dynamics by FRAP provided further insight into how differentiation reshapes chromatin engagement. Although the rate of fluorescence recovery was similar in mESCs and differentiated cells, suggesting that diffusional exchange was unchanged, the extent of recovery was significantly higher in differentiated cells, indicating a larger mobile fraction. This pattern suggests that differentiation reduces stable cohesin–chromatin residence while preserving rapid molecular turnover, consistent with either decreased lifetime of chromatin-bound cohesin or redistribution into a nucleoplasmic, chromatin-unengaged pool. These observations are consistent with decreased Sororin and ESCO1 levels, increased HDAC8 expression, and a net shift toward fewer stable cohesin–chromatin interactions. Rather than increasing rigidity by strengthening individual cohesin–chromatin binding events, differentiation appears to impose regulatory constraint on chromatin function while permitting greater cohesin mobility. Here, “regulatory constraint” refers to restriction of transcriptional flexibility rather than physical stiffening of chromosomes or direct alteration of long-range chromatin contacts. Together, the data support a model in which cohesin–chromatin interactions become less persistent at the molecular level during differentiation. Our findings place cohesin regulation within broader chromatin transitions that accompany lineage commitment. While pluripotent cells deploy cohesin to support architectural plasticity and long-range regulatory connectivity, differentiation is associated with reduced cohesin residence. By remodeling the balance between cohesin loading, stabilization, and release, cells reprogram chromatin mechanics during fate specification. This transition likely restricts lineage-inappropriate regulatory interactions and promotes the consolidation of lineage-specific gene-expression programs.

Our findings place cohesin regulatory proteins, rather than cohesin itself, at the center of chromatin reprogramming during lineage commitment. Differentiation is accompanied not by loss of the cohesin complex, but by changes in the abundance and activity of its S phase-independent regulators, including WAPL, ESCO1, and HDAC8, which together determine cohesin residence, stability, and functional output on chromatin. By modulating cohesin loading, stabilization, and release, these factors tune chromatin engagement and transcriptional changes as cells transition from pluripotency to lineage specification. These data emphasize that developmental regulation is mediated, in part, by the control of cohesin regulators and establish WAPL as a potential central node linking chromosome organization to transcriptional state during differentiation. We propose a model in which lineage commitment is driven, in part, by coordinated changes in cohesin regulators, which, in turn, tune chromatin engagement by cohesin without significant changes in cohesin abundance.

### LIMITATIONS OF THE STUDY

This study has several limitations. First, although the vermicelli phenotype was scored at the single-cell level by microscopy, transcriptional and protein-level measurements were performed in bulk, precluding resolution of cell-to-cell variability in gene expression or cohesin abundance. Second, the *in vitro* differentiation system used here yields a heterogeneous population rather than a uniform cardiomyocyte culture. While differentiated cultures robustly downregulated pluripotency markers such as OCT4 and formed contractile cell clusters and cardiomyocyte-associated gene expression, the precise cellular composition at later time points remains incompletely defined. It is therefore possible that some observed changes in cohesin regulators reflect lineage trajectories other than cardiomyocytes within the culture. Nevertheless, the consistent changes in WAPL, ESCO1, and other cohesin regulators across independent differentiation replicates indicate that regulation of cohesin dynamics is a robust feature of exit from pluripotency. Finally, this work examines a single differentiation trajectory and whether similar regulatory mechanisms act across other lineages or *in vivo* remains to be determined. Future single-cell and lineage-resolved studies will be required to fully define how cohesin regulation is coupled to cell fate transitions during development.

## Materials and Methods

### Mouse embryonic stem cell culture

The TC1 mouse embryonic stem cell line was a gift from Dr. Gerald Crabtree (Stanford University). TC1 mESCs were maintained in complete mESC medium (KnockOut DMEM (Gibco) supplemented with 7.5% ES-sure FBS (Omega Scientific), 7.5% KnockOut Serum Replacement (Gibco), 1X Penicillin/Streptamycin (Gibco), 1X GlutaMAX (Gibco), 1X MEM Non-Essential Amino Acids (Gibco), 1X Sodium Pyruvate (Gibco), 1:1000 β-mercaptoethanol (Gibco), and Leukemia Inhibitory Factor (LIF). Cells were grown feeder-free under standard cell culture conditions (37 °C, 5% CO₂), with media replaced daily. Cells were passaged every other day; upon reaching 80% confluence, cells were trypsinized with 0.25% trypsin-EDTA (Gibco) and resuspended in complete mESC medium to quench trypsin, centrifuged at 250 g for 4’ and resuspended in fresh complete mESC medium at 2 million cells/ 56cm^2^. Mouse embryonic fibroblasts, a gift from Dr. Gerald Crabtree, were treated with mitomycin C (10 μg/ml for 2-4 hours) and used for feeder layers as specified.

### Cell line construction

TC1-derived cell lines expressing degron-tagged alleles of WAPL or ESCO1 proteins were generated through CRISPR/Cas9 knock-in of a degron tag (FKBP12^F36V^) into the coding sequence of the C-terminal end of each protein. gBLOCKs containing linker, degron tag, and V5 protein tag were synthesized by IDT (Integrated DNA Technologies). Gene blocks containing homology arms (for WAPL, 497bp downstream and 540bp upstream; for ESCO1, 159bp downstream and 619bp upstream) were cloned into pBluescript II KS(+) plasmid (Stratagene) by isothermal assembly (Gibson et al., 2009), and sequences were confirmed by in-house long-read sequencing. To perform CRISPR knock-in, sgRNAs targeting Wapl or Esco1 genes near the stop codons were designed using the CHOP-CHOP web tool (chopchop.cbu.uib.no) and synthesized by IDT. Then, DNA oligos encoding sgRNAs were annealed and cloned into a plasmid backbone carrying the sgRNA scaffold and the Cas9 protein (pSpCas9(BB)-2A-Puro (PX459) V2.0; Addgene, #62988) (Ran et al., 2013). TC1 cells were transfected with the mixture of plasmids using a NEON electroporation system (Invitrogen). Briefly, 2e6 mESCs were mixed with 8 μg of plasmid carrying the gBLOCK and 4μg of plasmid encoding the gRNA. Cells were electroporated with 3 pulses (pulse voltage: 1.4V; pulse width: 10ms) and then plated on 6cm gelatin-coated plates with mitomycin-C-treated mouse embryonic fibroblasts (MEF) feeder cells. The next day, puromycin was added at 3 μg/mL and replaced each day for two additional days. Single colonies were picked, trypsinized, and plated in wells of a gelatin-coated 24-well plate with a MEF feeder layer. After reaching 80% confluency, cells were passaged and cultured until 100% confluency was achieved. Homozygous clones were then verified by sequencing of the PCR products and immunoblot analysis. WAPL or ESCO1 was degraded by the addition of 500nM dTAG^V^-1 (Tocris Biosciences) to the culture for 24h.

TC1-derived cells containing Halo-tagged RAD21 protein were made by CRISPR/Cas9 knock-in of a cassette encoding a linker peptide and HaloTag in frame with the *Rad21* coding sequence, followed by a stop codon, and then by a neomycin resistance gene flanked by Frt sequences (pEN313-Rad21-Halo-Frt-PGK-EM7-NeoR-bpA-Frt insert, Addgene #156431) (Nora et al., 2020). To perform the knock-in, a vector (pX330-EN1082_Rad21_STOP, Addgene #156450) (Nora et al., 2020) encoding Cas9 and sgRNA targeting Rad21 near the stop codon was used. Both constructs were introduced using the NEON electroporation system as above, and then plated on 6cm gelatin-coated plates with MEF feeder cells. The next day, neomycin selection was started (3μg/mL) and maintained for 5 days. After this selection, single colonies were picked, trypsinized, and plated on a gelatin-coated 24-well plate with a MEF feeder layer. After reaching 80% confluence, cells were passaged and subsequently cultured until 100% confluence. Genomic DNA was extracted from individual clones using Quick-Extract DNA solution (Biosearch Technologies) per manufacturer’s instructions, and homozygous clones were then confirmed by immunoblot. The FLP recombinase gene (Addgene #60662) (Xue et al., 2014) was introduced through electroporation, as above. Clones that expressed HaloTagged Rad21 and were sensitive to neomycin were picked for further experiments.

### Embryoid body (EB)-mediated cardiomyocyte differentiation

Upon reaching 70-90% confluence, cells were collected with trypsin and resuspended in EB medium (DMEM supplemented with 15% heat-inactivated FBS (Omega Scientific), Penicillin/Streptamycin (100U/ml), GlutaMAX, MEM Non-Essential Amino Acids, Sodium Pyruvate, 1:1000 β-mercaptoethanol (all Gibco), and 50 μg/mL L-ascorbic acid (STEMCELL Technologies)). To generate EBs, 20 μl droplets containing 1000 cells each were dispensed onto the inner surface of the lid of an untreated 150mm bacterial-grade Petri dish using a multi-channel pipette. The bottom of the Petri dish was filled with PBS to prevent drying, and the lid with droplets was carefully inverted back onto the dish. Plates were incubated in standard cell culture conditions (37°C, 5% CO₂) for 48 hours to allow EB formation. Then, each droplet containing a single EB was transferred into the well of a 24-well plate coated with 0.1% gelatin-coated (Stemcell Technologies) containing 1mL of fresh medium.

### Dissociation of differentiated cultures

On the desired day of differentiation, cultures were washed with Hank’s Balanced Salt Solution (HBSS) (Gibco) and incubated with trypsin for 5 minutes at 37 °C. Cells were detached by pipetting up and down in trypsin and subsequently transferred into Falcon tubes containing EB medium to neutralize the trypsin. After centrifugation at 250 × g for 5 minutes, the supernatant was aspirated, and the cell pellet was incubated with 1 mg/mL collagenase type II (Worthington Biochemical Corporation) in HBSS at 37 °C for 30 minutes. The suspension was triturated every 10 minutes to ensure dissociation into single cells, then quenched with EB medium, washed with 1× PBS. The collagenase step was omitted before day 6 of differentiation, at which point cells were dissociated and counted after trypsinization alone. For day 2 samples, EBs were collected individually, transferred into Falcon tubes pre-filled with EB medium, centrifuged at 250 × g for 5 minutes, then trypsinized and counted.

### EdU Incorporation assay

Cells were labeled by supplementing the media with 20 µM 5-ethynyl-2′-deoxyuridine (Thermo Fisher Scientific) for 30 minutes. The cells were washed once with PBS, trypsinized, and 1e^6^ cells were collected for analysis. After an additional PBS wash, cells were resuspended in a hypotonic lysis buffer (0.1% sodium citrate and 0.03% NP40) for 6 minutes on ice. Cells were then washed with 1% BSA in PBS and centrifuged at 1300 × g for 5 minutes. Cell pellets were resuspended in a labeling mix (PBS, 2 mM CuSO₄, 2 µM Alexa Fluor 647 azide (Life Technologies), and 50 mM ascorbic acid) and incubated for 75 min in the dark. Cells were washed once with PBS, then resuspended in a DNA staining solution containing 10 µg/mL propidium iodide and 100 µg/mL RNase A. Cell-cycle distributions were measured via flow cytometry (Beckman Coulter CytoFlex) and analyzed using FlowJo software (Tree Star, Inc.).

### Immunofluorescence staining

Cells grown on an 8-chamber cover glass-bottom dish (Lab-Tek) were washed with PBS three times. Cells were fixed in PBS containing 4% formaldehyde and 0.1% Triton X-100 for 15 minutes at room temperature. Cells were washed with PBS three times, and then blocked in antibody dilution buffer (AbDil: 20mM Tris pH=7.4, 150mM NaCl, 2% BSA, 0.1% Triton X-100, 0.1% NaN_3_) for 1 hour. Samples were incubated in the primary antibody diluted in AbDil overnight at 4°C. Cells were then washed three times (5 minutes per wash) in AbDil and incubated with secondary antibody for 1 hour at room temperature. Samples were washed in AbDil three times, 5 minutes each, counterstained with DAPI (Invitrogen), and washed again with PBS. Cells were kept in PBS for imaging on a C2 laser scanning confocal-equipped Nikon Eclipse Ti microscope.

### “Vermicelli” phenotype analysis

Cells were plated on a 6-well plate and treated with DMSO for 72 hours, or with dTAG^V^-1 for 72, 48, or 24 hours. Cells were trypsinized and replated 24 hours prior to analysis onto Lab-Tek Chambered Coverglass 8-well microscopy plates coated with Geltrex (Gibco). To degrade WAPL in cells that had already undergone differentiation, hanging drops were formed (as above) and then transferred to Lab-Tek chambered coverglass plates (1 EB per well), where they were further differentiated. On day 9 of differentiation, cells were treated with DMSO or dTAG^V^-1 for 24 hours. For vermicelli scoring, cells were immunostained with anti-SMC3 antibody and Alexa Fluor 568-labeled secondary antibody.

The percentage of “vermicelli” positive cells was blindly scored by counting 100 cells per condition per biological replicate.

### Fluorescence recovery after photobleaching (FRAP)

To study cohesin mobility, Fluorescence Recovery After Photobleaching (FRAP) was performed. Rad21-HaloTagged mESCs and differentiated cells were plated on Lab-Tek Chambered Coverglass 8-well microscopy plates coated with Geltrex (Gibco). 3.5×10^4^ mESCs or 5.0×104 differentiated cells per well were plated. After 48 hours, cells were incubated with HaloTag® Oregon Green® Ligand (Promega) for 30 minutes. Cells were then washed with fresh media and incubated with fresh media for 30 minutes to recover. Just before FRAP medium was swapped for Opti-MEM (Gibco), which doesn’t contain phenol red – a dye that can increase the background noise during fluorescence experiments. Plates were placed under the C2 confocal microscope (Nikon) and observed using a 60x objective and immersion oil. To visualize Oregon Green® Ligand, we used an excitation wavelength of 495 nm and an emission wavelength of 520 nm, which are the same as those for FITC. Part of the nuclei in 3 cells per technical replicate were photobleached at 70% laser power, and the cells were imaged for 26 minutes, with a new image acquired every 2 minutes. Images were analyzed with FIJI (Schindelin et al., 2012).

### Western blots

Whole cell extracts were prepared using 1x sample buffer or RIPA buffer (50 mM Tris pH 7.8, 150 mM NaCl, 1% NP-40, 1% SDS and 0.1% sodium deoxycholate), 1mM DTT, protease inhibitors (leupeptin, pepstatin, and chymostatin (10μg/ml each; EMD Millipore) and 1.25 U/μl Benzonase (Millipore). Protein samples were resolved on 7–15% Tris-glycine gradient polyacrylamide gels (homemade) or 4-18% Bis-Tris protein gels (Invitrogen) and transferred to nitrocellulose membranes using Trans Blot Turbo Transfer System (Bio-Rad). Membranes were blocked in 5% milk in Tris-buffered saline (TBST: 20mM Tris pH 7.5, 150mM NaCl, 0.05% Tween 20, or Intercept Protein-Free Blocking Buffer (Li-Cor) for 1h, then incubated with primary antibody diluted in 5% milk in TBST and 0.1% NaN_3_ or Intercept T20 Antibody Diluent (Li-Cor) overnight at 4°C. Membranes were washed three times in TBST and incubated with a secondary antibody in TBST containing 5% milk or in T20 Antibody Diluent (Li-Cor) with 0.01% SDS for 45 min. After the membranes were washed three times in TBST and once in TBS, bands were detected with chemiluminescence substrate (Azure Biosystems) and imaged using C600 Imager (Azure) or Odyssey DLx imaging system (Li-Cor). Protein band intensities were quantified using Fiji/ImageJ and normalized to β-tubulin, or to histone H2B in the case of Cyclin B1. All uncropped western blots used in the figures are provided in a blot transparency file.

### RNA-sequencing

For transcriptional analysis, TC1 cells were dissociated either before differentiation or on day 10 of the protocol. Total RNA was extracted from a single-cell suspension using TRI Reagent (Sigma-Aldrich) according to the manufacturer’s instructions. Library preparation was done using SMART-Seq Total RNA High Input (RiboGone Mammalian). Libraries will be run on a Tapestation, quantified using a Qubit fluorometer, and sequenced at the OMRF Clinical Genomics Core on a NovaseqX (Illumina). Quality control of samples was performed with fastqc (v0.12.1) (Andrews). Reads were pseudo-aligned to the mouse genome (index: V1 mouse_index_standard) using Kallisto V1 (Bray et al., 2016). Differential expression analysis of read count files (.tsv) was performed in DESeq2 (Love et al., 2014a; Love et al., 2014b). Differentially expressed gene lists used cutoffs of (adj p-value < 0.05) and (log2FoldChange > 0.585 | log2FoldChange < −0.585). Log2-fold change (log2FC) and lcf standard error (SE) were calculated in DESeq2.

### RT-qPCR analysis

RNA was extracted from 2e6 cells using TRI Reagent (Sigma Aldrich), followed by chloroform extraction, ethanol precipitation, and resuspension in 30 μl of nuclease-free water. cDNA was synthesized by reverse transcription using 1μg of RNA and the SensiFAST cDNA Synthesis Kit (Meridian) according to the manufacturer’s instructions. The cDNA was diluted 1:4 with water, and in each qPCR reaction, 2 μL of diluted cDNA was amplified using 2X qMAX SYBR Green low ROX reagent (Accuris) on a QuantStudio 3 Flex system (Applied Biosystems).

### Cell viability assays

TC-1 cells and their derivatives were plated on a 96-well plate (500 cells/well, in 100 μL media) and treated daily with media containing DMSO or dTAG^V^-1 for 0, 1, 2, 3, 4, or 5 days. For the final 10 hours of culture, 10 μl of CellTiter-Blue (Promega) was added to each well. After 10 hours, the supernatant was transferred to a fresh 96-well plate and analyzed in the GloMax Plate Reader (Promega) with an excitation wavelength of 544nm and an emission wavelength of 590nm.

Cells were cultured on Lab-Tek Chambered Coverglass 8-well microscopy plates pre-coated with Geltrex (Gibco). Undifferentiated cells (day 0) and cells differentiated for 10 days were used for the assay. Prior to staining, culture medium was removed and cells were washed once with Hanks’ Balanced Salt Solution (HBSS). Cell viability was assessed using the LIVE/DEAD® Reduced Biohazard Viability/Cytotoxicity Kit #1 (L-7013; Molecular Probes). SYTO 10 green fluorescent nucleic acid stain and DEAD Red nucleic acid stain were mixed and diluted in HBSS according to the manufacturer’s instructions. Cells were incubated with the dye mixture for 15 minutes at room temperature in the dark. Following incubation, cells were washed once with HBSS and fixed with freshly prepared 4% glutaraldehyde in HBSS for 1 hour at room temperature. Cells were imaged using a Nikon Eclipse Ti microscope equipped with a C2 laser scanning confocal system.

### Statistical analyses

All statistical analyses in this work were performed in GraphPad Prism 10 (GraphPad Software, Inc.). All experiments were performed in at least three biological replicates. A Student’s t-test was performed to compare two samples. One- or two-way ANOVA was used for multiple comparisons, followed by Dunnett or Šídák’s post-hoc tests. When heterogeneity of variance was detected, the Brown–Forsythe test was applied, and appropriate post-hoc corrections were used. For FRAP analysis a linear mixed model with time and cell type as fixed effects was used. Homoskedasticity and linearity were confirmed. Normality assumptions were confirmed with QQPlots. The model considered time, cell type and the interaction between the two.

### Antibodies used in this study

| Antigen | Antibody source | Number | Dilution (IF/IB) |
| --- | --- | --- | --- |
| β-tubulin | Development Studies<br>Hybridoma Bank | E7 | 1:4000 |
| Cyclin B1 | Santa Cruz Biotechnology | SC-752 | 1:1000 |
| Donkey α-mouse<br>(HRP conjugated) | Jackson ImmunoResearch | 715035151 | 1:5000 |
| Esco1 | Lab made | PRF&L35698 | 1:2000 |
| Goat α-rabbit (HRP<br>conjugated) | Thermo Fisher Scientific | 31460 | 1:5000 |
| H2B | Cell Signaling Technology | 53H3 | 1:1000 |
| HDAC8 | Santa Cruz Biotechnology | SC-17778 | 1:1000 |
| Rad21 | Thermo Fisher Scientific | PA5-120663 | 1:1000 |
| Smc3 | Lab made | OMRF160 | 1:1000 |
| Sororin | Lab made | OMRF167 | 1:3000 |
| Wapl | Cell Signaling Technology | D9J1U | 1:1000 |
| Goat α-rabbit AF568 | Invitrogen | A-11036 | 1:1000 |

### Plasmids used in this study

#### Esco1_dTAG

**Plasmids used in this study****Esco1\_dTAG**
| Name | Description | Parent plasmid(s) |
| --- | --- | --- |
| pKT4 | pJK307 (Esco1_gRNA_10) | pSpCas9(BB)-2A-Puro (PX459) V2.0 #62988 |
| pKT5 | pJK307 (Esco1_gRNA_26) | pSpCas9(BB)-2A-Puro (PX459) V2.0 #62988 |
| pKT6 | pJK307 (Esco1_gRNA_177) | pSpCas9(BB)-2A-Puro (PX459) V2.0 #62988 |
| pKT8 | pJK308 (Esco1_dTAG_homology) | pBluescript-II KS+ |

#### Wapl_dTAG

| Name | Description | Parent plasmid(s) |
| --- | --- | --- |
| pKT1 | pJK307 (Wapl_gRNA_153) | pSpCas9(BB)-2A-Puro (PX459) V2.0 #62988 |
| pKT2 | pJK307 (Wapl_gRNA_221) | pSpCas9(BB)-2A-Puro (PX459) V2.0 #62988 |
| pKT3 | pJK307 (Wapl_gRNA_26) | pSpCas9(BB)-2A-Puro (PX459) V2.0 #62988 |
| pKT7 | PJK308 (Wapl_dTAG_homology) | Bluescript-II KS+ |

pSpCas9(BB)-2A-Puro (PX459) V2.0 was a gift from Feng Zhang (Addgene plasmid # 62988 ; http://n2t.net/addgene:62988 ; RRID:Addgene_62988). pEN313 - Rad21-Halo-Frt-PGK-EM7-NeoR-bpA-Frt targeting was a gift from Elphege Nora (Addgene plasmid # 156431 ; http://n2t.net/addgene:156431 ; RRID:Addgene_156431). pX330-EN1082_Rad21_STOP was a gift from Elphege Nora (Addgene plasmid # 156450 ; http://n2t.net/addgene:156450 ; RRID:Addgene_156450). pCAG-Flpo was a gift from Massimo Scanziani (Addgene plasmid # 60662 ; http://n2t.net/addgene:60662 ; RRID:Addgene_60662)

### Primers used in this study

#### Esco1 manipulation and analysis

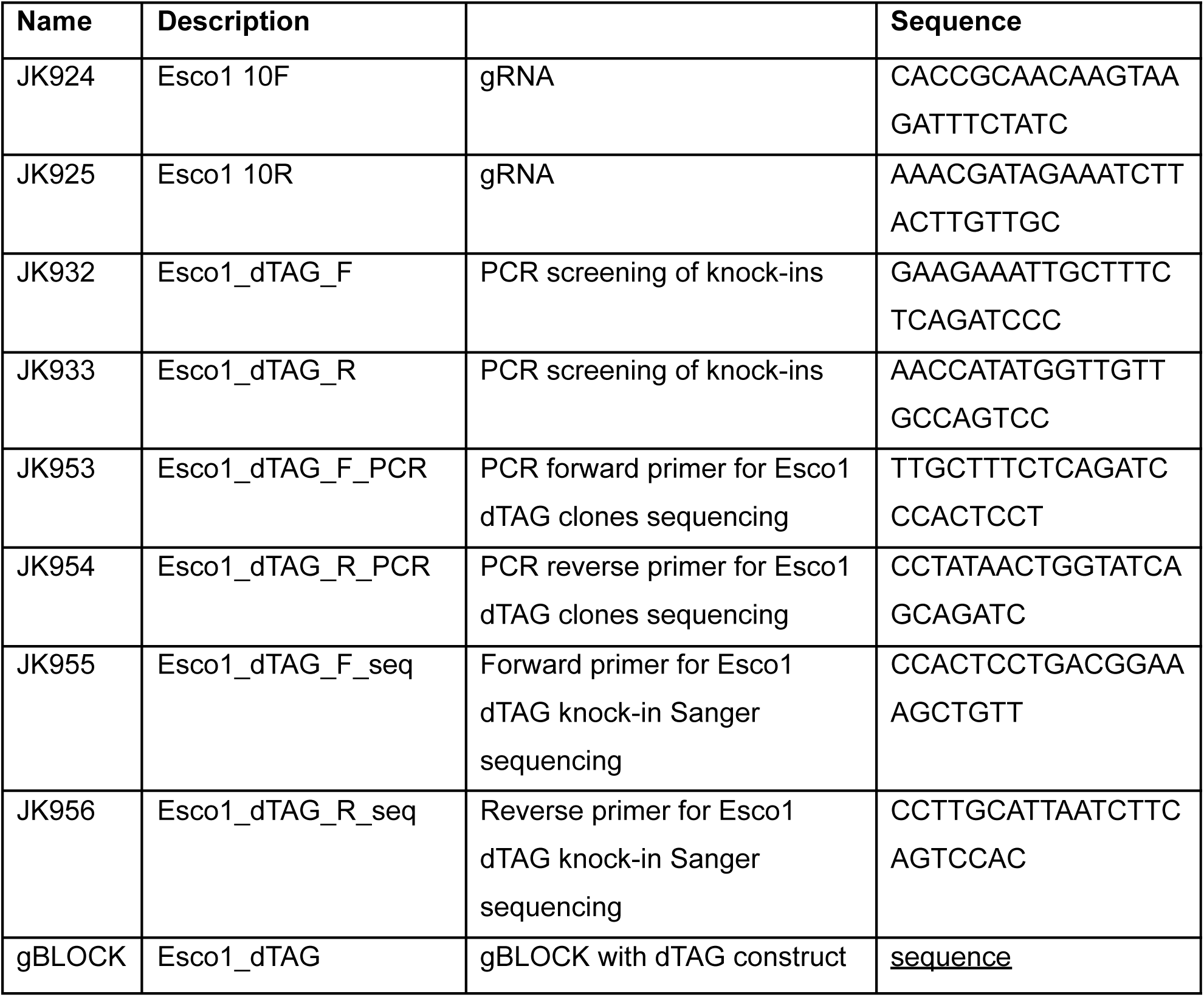

#### Wapl manipulation and analysis

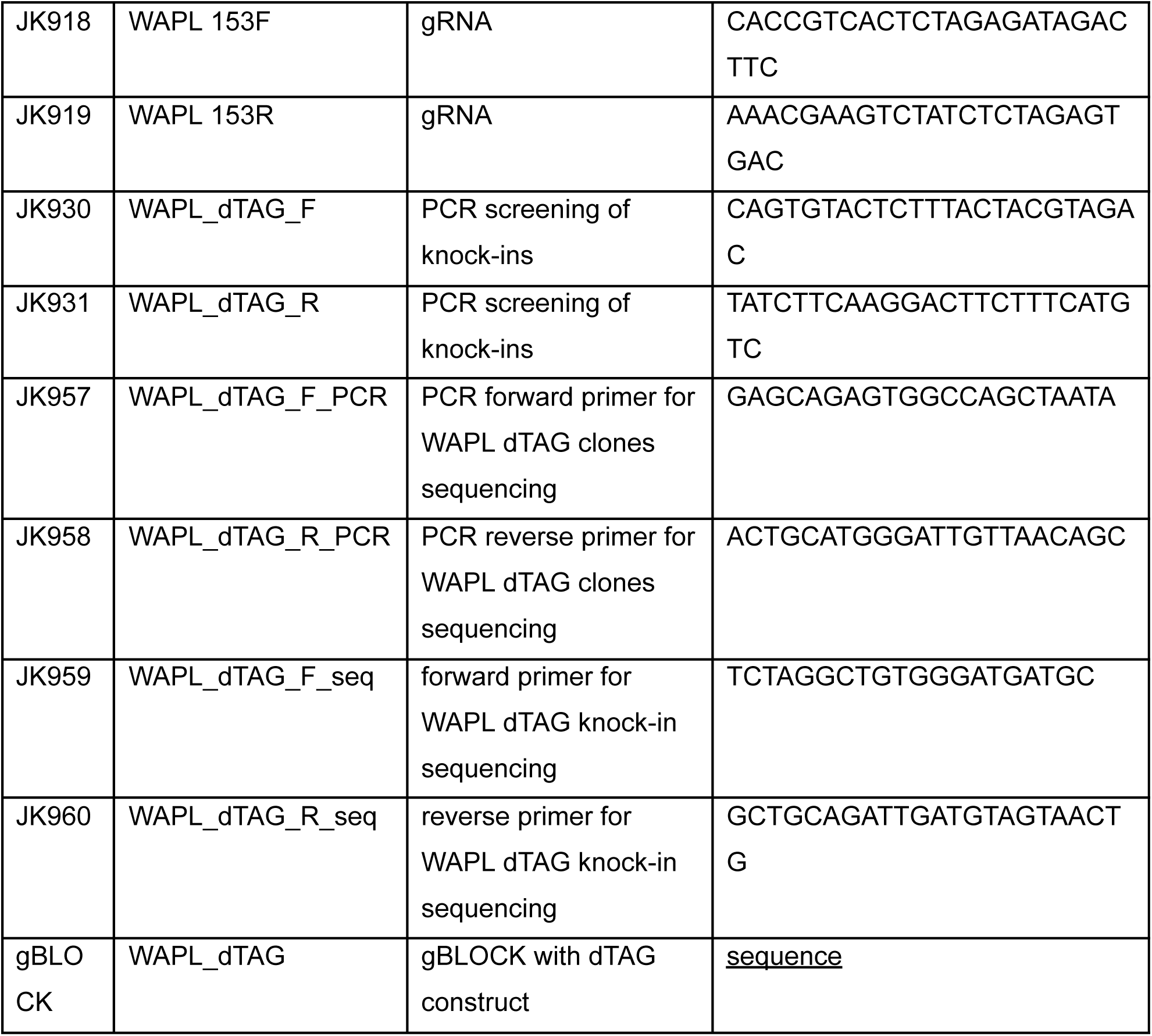

#### qPCR primers

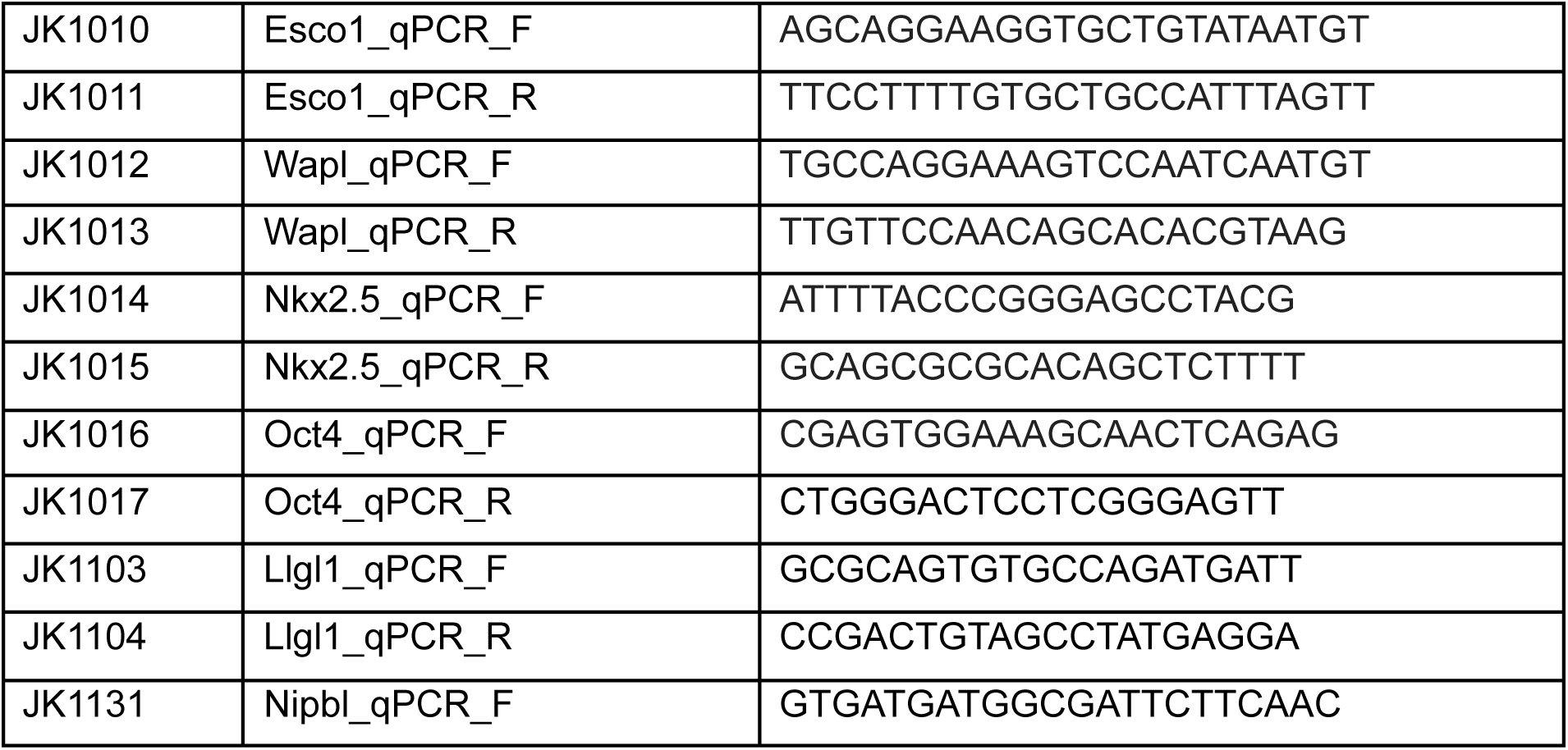
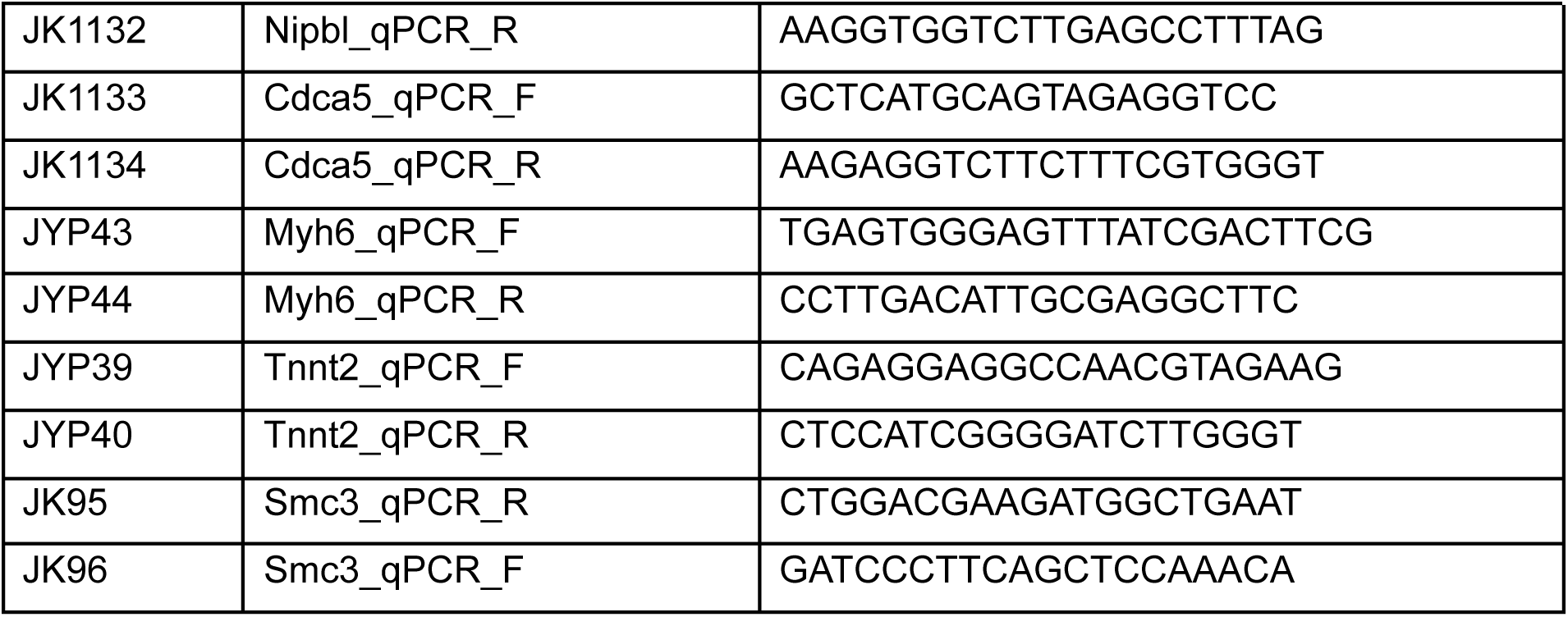

Detailed plasmid sequences and maps gladly provided upon request.

## ACKNOWLEDGEMENTS

Funding was provided by a Presbyterian Health Foundation collaborative grant (J.G.K. and S.R.), 5R35GM154839 (J.K.), and R35GM149343 (to S.R.) Data processing and analysis were supported by the OMRF Center for Biomedical Data Sciences.

---

**Figure S1.**
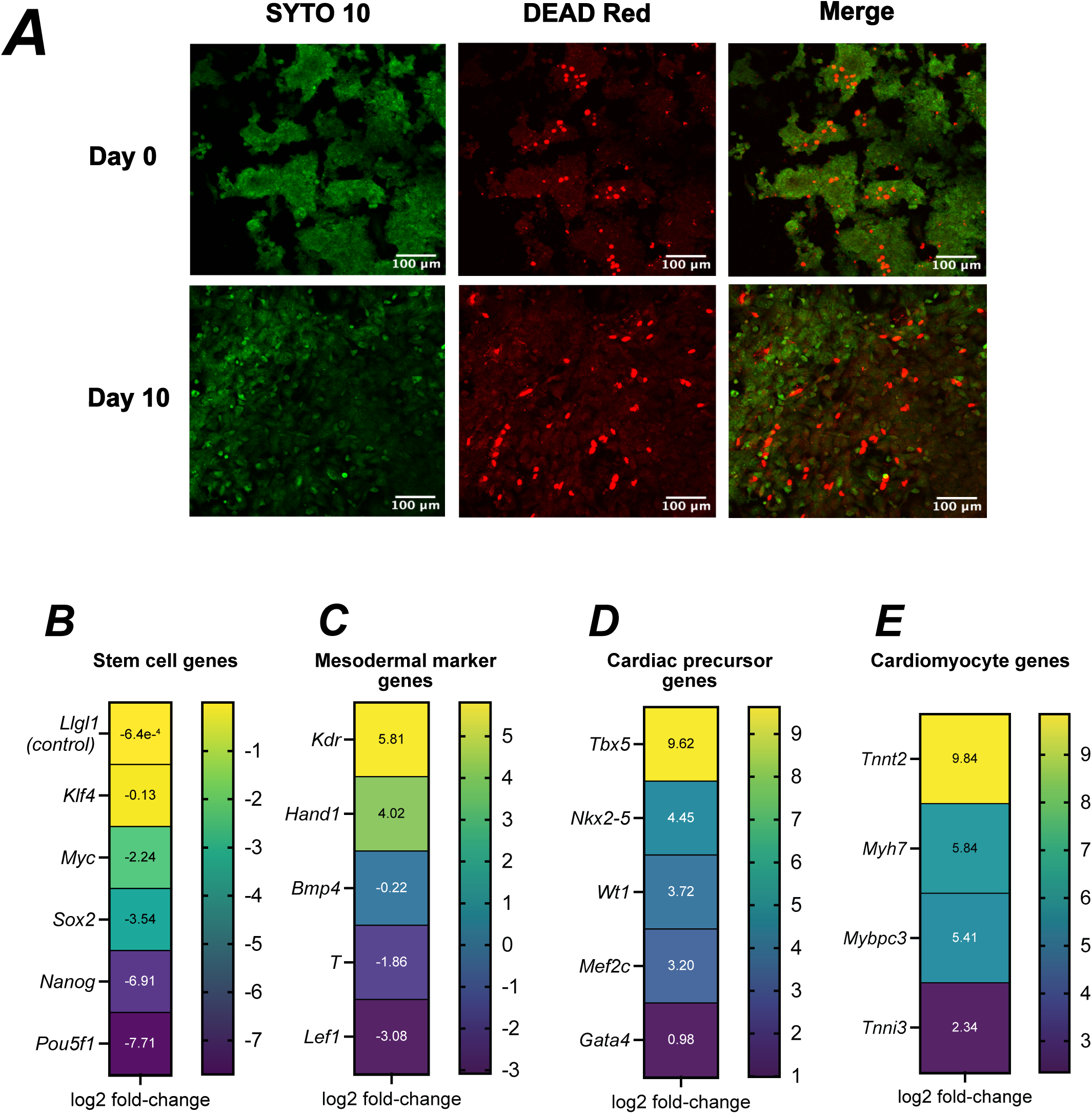
Differentiation of mESCs. **A. Cell death by SYTO10/DEAD Red**. Images from undifferentiated cells (day 0) and differentiated cells (day 10) using SYTO 10 to stain DNA in the nuclei of all cells and DEAD Red to stain DNA of dead cells. Scale bar = 100 µm. **B-D. Gene expression changes during differentiation obtained from bulk RNAseq. B.** Heat map indicating changes in expression of stem cell genes. **C.** Heat map indicating changes in expression of mesodermal genes. **D.** Heat map indicating changes in expression of cardiac precursor genes. **E.** Heat map indicating changes in expression of cardiomyocyte-specific genes. Values are log2 fold-change with different scaling for B-E.

**Figure S2.**
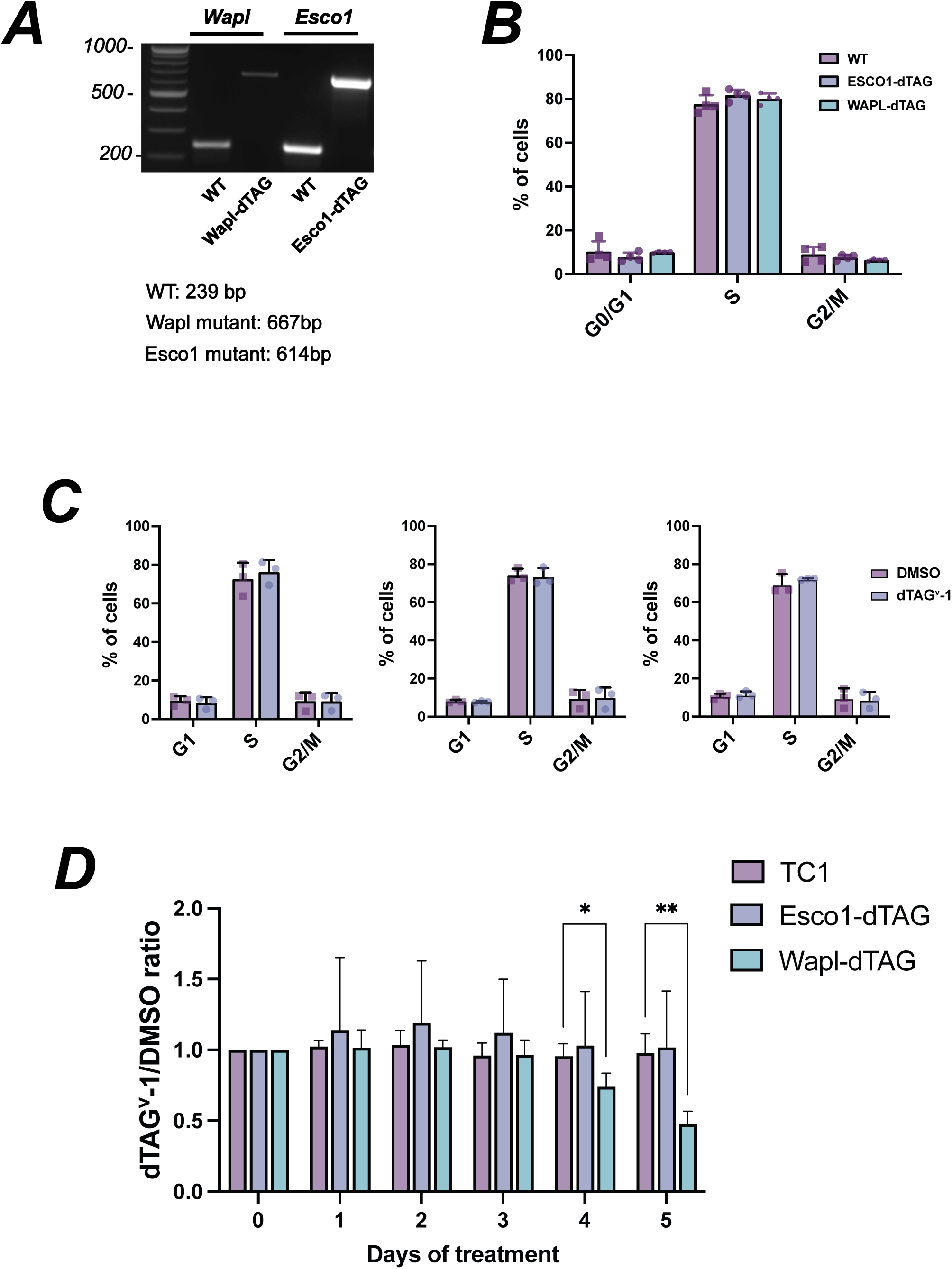
Characterization of WAPL-dTAG and ESCO1-dTAG mESC lines. **A. Genotyping CRISPR incorporation of dTAG fusions.** Ethidium bromide stained PCR products generated using gene-specific primers indicated in Figure 4A (green arrows). **B. Cell cycle analysis.** Percentages of cells within each gate for G0/G1, S, and G2/M phases in WT parental and ESCO1-dTAG or WAPL-dTAG untreated cells shows no effect on the cell cycle (n=4). **C. Cell cycle analysis.** Percentages of cells within each gate for G0/G1, S, and G2/M phases after 24 hours of dTag^V^-1 treatment (n=3). **D. CellTiter-Blue cell viability assay**. Ratio of fluorescence of dTag^V^-1/DMSO treated cells at 1, 2, 3, 4, and 5 days normalized to untreated cells (day 0). Statistically significant growth defects in the WAPL-dTAG line were detected starting at day 4. n=5. * = p <0.05, ** = p <0.01 by 2-way ANOVA with Dunnett’s multiple comparison test.

***Blot Transparency - Figure 3A***

***Azure Imager***

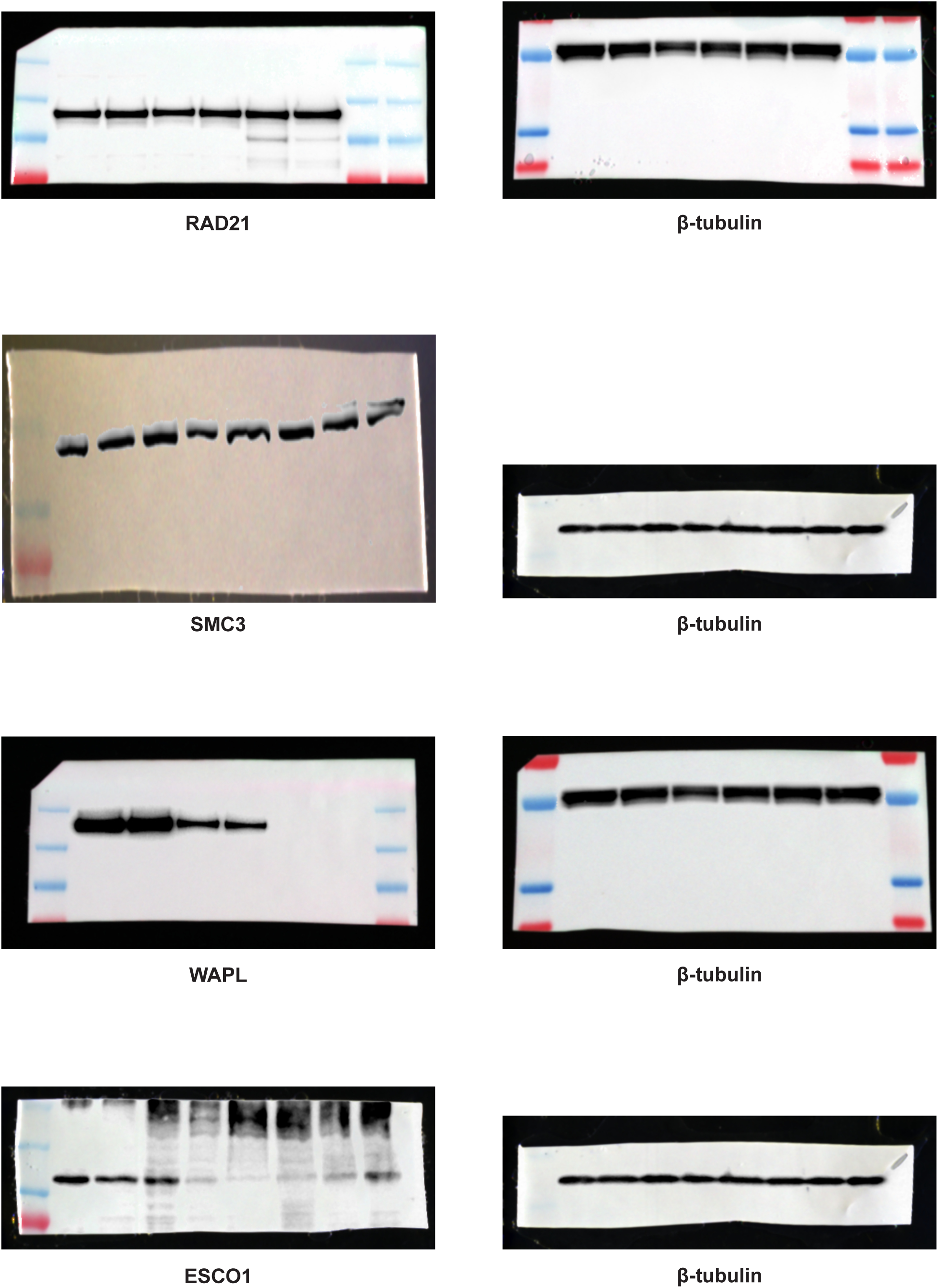
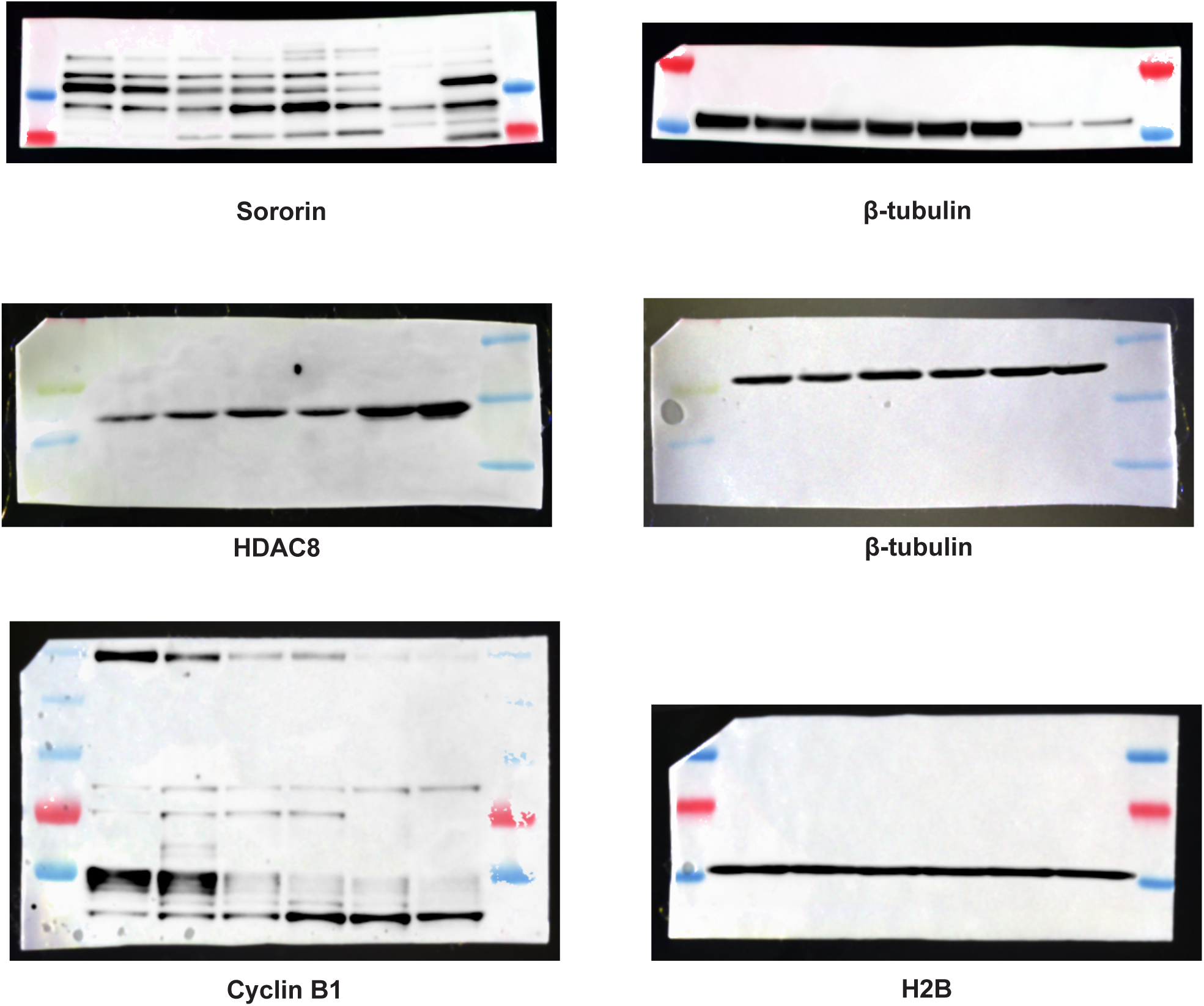

***Blot Transparency - Figure 4B***

***Azure Imager***

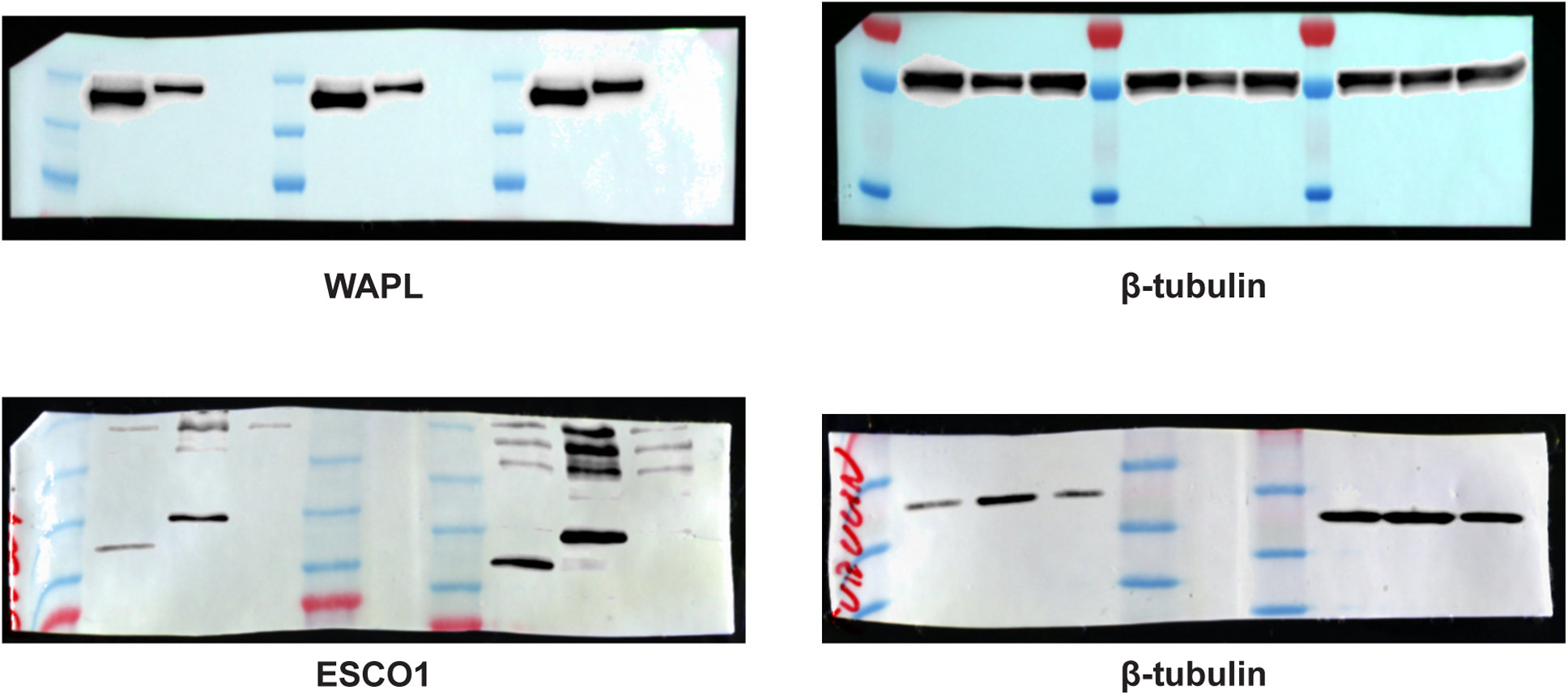

***Blot Transparency - Figure 6A***

***LiCor Imager***

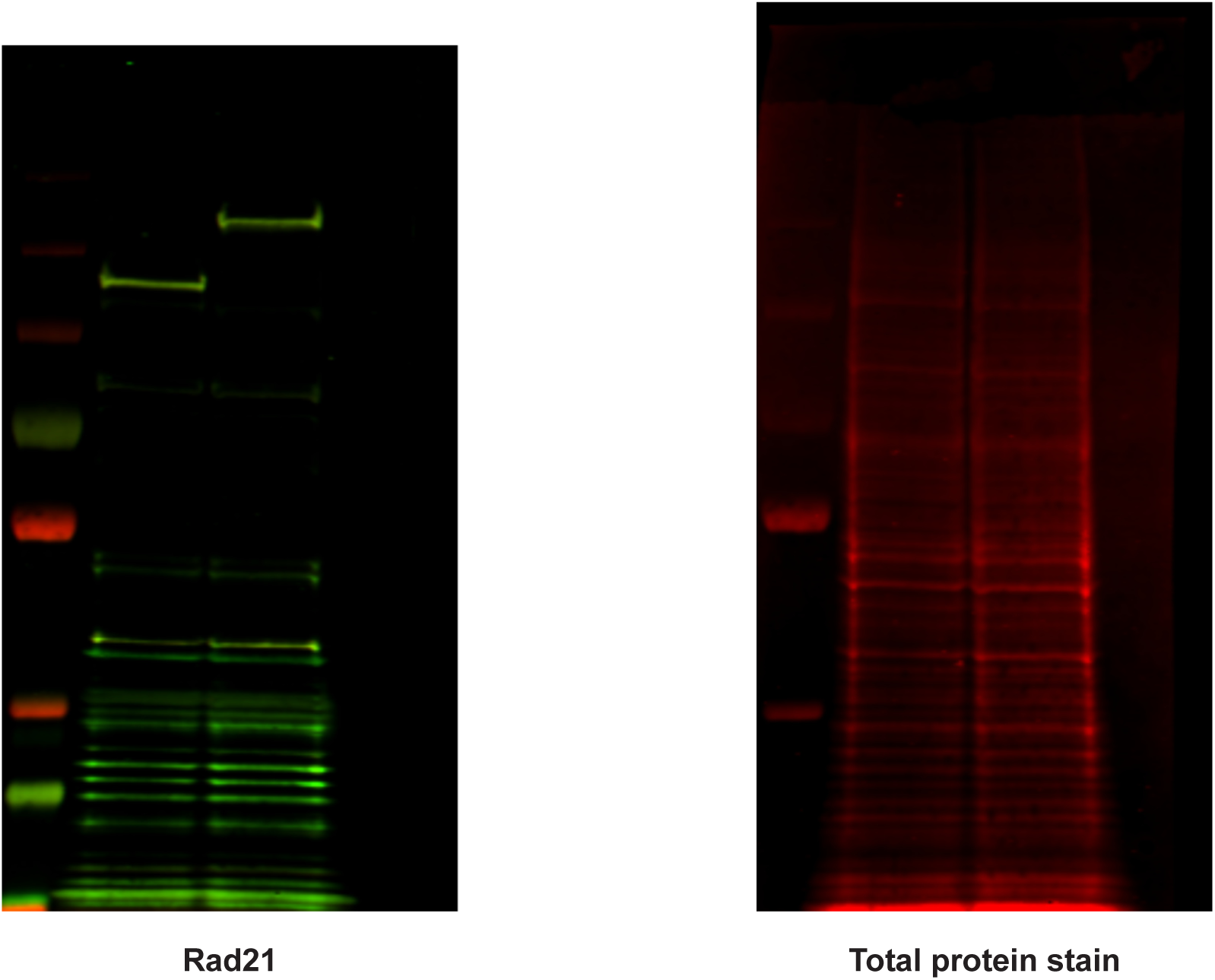

## REFERENCES

Andrews, S. FastQC: A quality control analysis tool for high throughput sequencing data. Github.

Bender, D., Da Silva, E. M. L., Chen, J., Poss, A., Gawey, L., Rulon, Z. and Rankin, S. (2020). Multivalent interaction of ESCO2 with the replication machinery is required for sister chromatid cohesion in vertebrates. Proc. Natl. Acad. Sci. U. S. A. 117, 1081–1089.

Bonev, B. and Cavalli, G. (2016). Organization and function of the 3D genome. Nat. Rev. Genet. 17, 661–678.

Bray, N. L., Pimentel, H., Melsted, P. and Pachter, L. (2016). Near-optimal probabilistic RNA-seq quantification. Nat. Biotechnol. 34, 525–527.

Bryan, A. F., Justice, M., Stutzman, A. V., McKay, D. J. and Dowen, J. M. (2025). Cohesin stabilization at promoters and enhancers by common transcription factors and chromatin regulators. Epigenetics Chromatin 18, 33.

Calder, A., Roth-Albin, I., Bhatia, S., Pilquil, C., Lee, J. H., Bhatia, M., Levadoux-Martin, M., McNicol, J., Russell, J., Collins, T., et al. (2013). Lengthened G1 phase indicates differentiation status in human embryonic stem cells. Stem Cells Dev. 22, 279–295.

Chen, H., Chen, J., Zhao, L., Song, W., Xuan, Z., Chen, J., Li, Z., Song, G., Hong, L., Song, P., et al. (2019). CDCA5, transcribed by E2F1, promotes oncogenesis by enhancing cell proliferation and inhibiting apoptosis via the AKT pathway in hepatocellular carcinoma. J. Cancer 10, 1846–1854.

Coronado, D., Godet, M., Bourillot, P.-Y., Tapponnier, Y., Bernat, A., Petit, M., Afanassieff, M., Markossian, S., Malashicheva, A., Iacone, R., et al. (2013). A short G1 phase is an intrinsic determinant of naïve embryonic stem cell pluripotency. Stem Cell Res. 10, 118–131.

Cuadrado, A., Giménez-Llorente, D., De Koninck, M., Ruiz-Torres, M., Kojic, A., Rodríguez-Corsino, M. and Losada, A. (2022). Contribution of variant subunits and associated factors to genome-wide distribution and dynamics of cohesin. Epigenetics Chromatin 15, 37.

Davidson, I. F., Bauer, B., Goetz, D., Tang, W., Wutz, G. and Peters, J.-M. (2019). DNA loop extrusion by human cohesin. Science 366, 1338–1345.

Deardorff, M. A., Bando, M., Nakato, R., Watrin, E., Itoh, T., Minamino, M., Saitoh, K., Komata, M., Katou, Y., Clark, D., et al. (2012). HDAC8 mutations in Cornelia de Lange syndrome affect the cohesin acetylation cycle. Nature 489, 313–317.

De Koninck, M., Lapi, E., Badía-Careaga, C., Cossío, I., Giménez-Llorente, D., Rodríguez-Corsino, M., Andrada, E., Hidalgo, A., Manzanares, M., Real, F. X., et al. (2020). Essential roles of cohesin STAG2 in mouse embryonic development and adult tissue homeostasis. Cell Rep. 32, 108014.

Dowen, J. M., Bilodeau, S., Orlando, D. A., Hübner, M. R., Abraham, B. J., Spector, D. L. and Young, R. A. (2013). Multiple structural maintenance of chromosome complexes at transcriptional regulatory elements. Stem Cell Reports 1, 371–378.

Dowen, J. M., Fan, Z. P., Hnisz, D., Ren, G., Abraham, B. J., Zhang, L. N., Weintraub, A. S., Schujiers, J., Lee, T. I., Zhao, K., et al. (2014). Control of cell identity genes occurs in insulated neighborhoods in mammalian chromosomes. Cell 159, 374–387.

Garcia, P., Fernandez-Hernandez, R., Cuadrado, A., Coca, I., Gomez, A., Maqueda, M., Latorre-Pellicer, A., Puisac, B., Ramos, F. J., Sandoval, J., et al. (2021). Disruption of NIPBL/Scc2 in Cornelia de Lange Syndrome provokes cohesin genome-wide redistribution with an impact in the transcriptome. Nat. Commun. 12, 4551.

Gerlich, D., Koch, B., Dupeux, F., Peters, J.-M. and Ellenberg, J. (2006). Live-cell imaging reveals a stable cohesin-chromatin interaction after but not before DNA replication. Curr. Biol. 16, 1571–1578.

Gibson, D. G., Young, L., Chuang, R.-Y., Venter, J. C., Hutchison, C. A., 3rd and Smith, H. O. (2009). Enzymatic assembly of DNA molecules up to several hundred kilobases. Nat. Methods 6, 343–345.

Guacci, V., Chatterjee, F., Robison, B. and Koshland, D. E. (2019). Communication between distinct subunit interfaces of the cohesin complex promotes its topological entrapment of DNA. eLife 8,.

Haarhuis, J. H. I., Elbatsh, A. M. O., van den Broek, B., Camps, D., Erkan, H., Jalink, K., Medema, R. H. and Rowland, B. D. (2013). WAPL-mediated removal of cohesin protects against segregation errors and aneuploidy. Curr. Biol. 23, 2071–2077.

Haarhuis, J. H. I., Elbatsh, A. M. O. and Rowland, B. D. (2014). Cohesin and its regulation: on the logic of X-shaped chromosomes. Dev. Cell 31, 7–18.

Hansen, A. S., Pustova, I., Cattoglio, C., Tjian, R. and Darzacq, X. (2017). CTCF and cohesin regulate chromatin loop stability with distinct dynamics. Elife 6,.

Heuvelmans, L., Mostert, D., Spanò, G., Stoll, M. and De Windt, L. J. (2025). GATA4: orchestrating cardiac development and beyond. Cardiovasc. Res. 121, 2476–2483.

Hota, S. K., Rao, K. S., Blair, A. P., Khalilimeybodi, A., Hu, K. M., Thomas, R., So, K., Kameswaran, V., Xu, J., Polacco, B. J., et al. (2022). Brahma safeguards canalization of cardiac mesoderm differentiation. Nature 602, 129–134.

Hsieh, T.-H. S., Cattoglio, C., Slobodyanyuk, E., Hansen, A. S., Darzacq, X. and Tjian, R. (2022). Enhancer-promoter interactions and transcription are largely maintained upon acute loss of CTCF, cohesin, WAPL or YY1. Nat. Genet. 54, 1919–1932.

Kueng, S., Hegemann, B., Peters, B. H., Lipp, J. J., Schleiffer, A., Mechtler, K. and Peters, J.-M. (2006). Wapl controls the dynamic association of cohesin with chromatin. Cell 127, 955–967.

Lafont, A. L., Song, J. and Rankin, S. (2010). Sororin cooperates with the acetyltransferase Eco2 to ensure DNA replication-dependent sister chromatid cohesion. Proc. Natl. Acad. Sci. U. S. A. 107, 20364–20369.

Liu, L., Michowski, W., Kolodziejczyk, A. and Sicinski, P. (2019). The cell cycle in stem cell proliferation, pluripotency and differentiation. Nat. Cell Biol. 21, 1060–1067.

Liu, N. Q., Magnitov, M., Schijns, M. M. G. A., van Schaik, T., Teunissen, H., van Steensel, B. and de Wit, E. (2025). Extrusion fountains are restricted by WAPL-dependent cohesin release and CTCF barriers. Nucleic Acids Res. 53,.

Love, M., Anders, S. and Huber, W. (2014a). Differential analysis of count data--the DESeq2 package. Genome Biol. 15, 550.

Love, M. I., Huber, W. and Anders, S. (2014b). Moderated estimation of fold change and dispersion for RNA-seq data with DESeq2. Genome Biol. 15, 550.

Lynch, A. T., Mazzotta, S. and Hoppler, S. (2018). Cardiomyocyte differentiation from mouse embryonic stem cells. Methods Mol. Biol. 1816, 55–66.

Meluzzi, D. and Arya, G. (2020). Computational approaches for inferring 3D conformations of chromatin from chromosome conformation capture data. Methods 181-182, 24–34.

Mfarej, M. G., Hyland, C. A., Sanchez, A. C., Falk, M. M., Iovine, M. K. and Skibbens, R. V. (2023). Cohesin: an emerging master regulator at the heart of cardiac development. Mol. Biol. Cell 34, rs2.

Nabet, B., Ferguson, F. M., Seong, B. K. A., Kuljanin, M., Leggett, A. L., Mohardt, M. L., Robichaud, A., Conway, A. S., Buckley, D. L., Mancias, J. D., et al. (2020). Rapid and direct control of target protein levels with VHL-recruiting dTAG molecules. Nat. Commun. 11, 4687.

Niwa, H., Miyazaki, J. and Smith, A. G. (2000). Quantitative expression of Oct-3/4 defines differentiation, dedifferentiation or self-renewal of ES cells. Nat. Genet. 24, 372–376.

Nora, E. P., Caccianini, L., Fudenberg, G., So, K., Kameswaran, V., Nagle, A., Uebersohn, A., Hajj, B., Saux, A. L., Coulon, A., et al. (2020). Molecular basis of CTCF binding polarity in genome folding. Nat. Commun. 11, 5612.

Ran, F. A., Hsu, P. D., Wright, J., Agarwala, V., Scott, D. A. and Zhang, F. (2013). Genome engineering using the CRISPR-Cas9 system. Nat. Protoc. 8, 2281–2308.

Rankin, S., Ayad, N. G. and Kirschner, M. W. (2005). Sororin, a substrate of the anaphase-promoting complex, is required for sister chromatid cohesion in vertebrates. Mol. Cell 18, 185–200.

Rao, S. S. P., Huang, S.-C., Glenn St Hilaire, B., Engreitz, J. M., Perez, E. M., Kieffer-Kwon, K.-R., Sanborn, A. L., Johnstone, S. E., Bascom, G. D., Bochkov, I. D., et al. (2017). Cohesin loss eliminates all loop domains. Cell 171, 305–320.e24.

Rhodes, J. D. P., Haarhuis, J. H. I., Grimm, J. B., Rowland, B. D., Lavis, L. D. and Nasmyth, K. A. (2017). Cohesin can remain associated with chromosomes during DNA replication. Cell Rep. 20, 2749–2755.

Rittenhouse, N. L. and Dowen, J. M. (2024). Cohesin regulation and roles in chromosome structure and function. Curr. Opin. Genet. Dev. 85, 102159.

Samejima, K., Gibcus, J. H., Abraham, S., Cisneros-Soberanis, F., Samejima, I., Beckett, A. J., Pučeková, N., Abad, M. A., Medina-Pritchard, B., Paulson, J. R., et al. (2024). Rules of engagement for condensins and cohesins guide mitotic chromosome formation. bioRxivorg.

Sansam, C. G., Pietrzak, K., Majchrzycka, B., Kerlin, M. A., Chen, J., Rankin, S. and Sansam, C. L. (2018). A mechanism for epigenetic control of DNA replication. Genes Dev. 32, 224–229.

Schindelin, J., Arganda-Carreras, I., Frise, E., Kaynig, V., Longair, M., Pietzsch, T., Preibisch, S., Rueden, C., Saalfeld, S., Schmid, B., et al. (2012). Fiji: an open-source platform for biological-image analysis. Nat. Methods 9, 676–682.

Sehnert, A. J., Huq, A., Weinstein, B. M., Walker, C., Fishman, M. and Stainier, D. Y. R. (2002). Cardiac troponin T is essential in sarcomere assembly and cardiac contractility. Nat. Genet. 31, 106–110.

Seitan, V. C., Faure, A. J., Zhan, Y., McCord, R. P., Lajoie, B. R., Ing-Simmons, E., Lenhard, B., Giorgetti, L., Heard, E., Fisher, A. G., et al. (2013). Cohesin-based chromatin interactions enable regulated gene expression within preexisting architectural compartments. Genome Res. 23, 2066–2077.

Solé-Ferran, M. and Losada, A. (2025). Cohesin in 3D: development, differentiation, and disease. Genes Dev. 39, 679–696.

Srinivasan, M., Fumasoni, M., Petela, N. J., Murray, A. and Nasmyth, K. A. (2020). Cohesion is established during DNA replication utilising chromosome associated cohesin rings as well as those loaded de novo onto nascent DNAs. Elife 9,.

Tedeschi, A., Wutz, G., Huet, S., Jaritz, M., Wuensche, A., Schirghuber, E., Davidson, I. F., Tang, W., Cisneros, D. A., Bhaskara, V., et al. (2013). Wapl is an essential regulator of chromatin structure and chromosome segregation. Nature 501, 564–568.

Veevers, J., Farah, E. N., Corselli, M., Witty, A. D., Palomares, K., Vidal, J. G., Emre, N., Carson, C. T., Ouyang, K., Liu, C., et al. (2018). Cell-surface marker signature for enrichment of ventricular cardiomyocytes derived from human embryonic stem cells. Stem Cell Reports 11, 828–841.

Waizenegger, I. C., Hauf, S., Meinke, A. and Peters, J. M. (2000). Two distinct pathways remove mammalian cohesin from chromosome arms in prophase and from centromeres in anaphase. Cell 103, 399–410.

Walsh, K. and Perlman, H. (1997). Cell cycle exit upon myogenic differentiation. Curr. Opin. Genet. Dev. 7, 597–602.

Wamstad, J. A., Alexander, J. M., Truty, R. M., Shrikumar, A., Li, F., Eilertson, K. E., Ding, H., Wylie, J. N., Pico, A. R., Capra, J. A., et al. (2012). Dynamic and coordinated epigenetic regulation of developmental transitions in the cardiac lineage. Cell 151, 206–220.

Watrin, E., Kaiser, F. J. and Wendt, K. S. (2016). Gene regulation and chromatin organization: relevance of cohesin mutations to human disease. Curr. Opin. Genet. Dev. 37, 59–66.

Wutz, G., Ladurner, R., St Hilaire, B. G., Stocsits, R. R., Nagasaka, K., Pignard, B., Sanborn, A., Tang, W., Várnai, C., Ivanov, M. P., et al. (2020). ESCO1 and CTCF enable formation of long chromatin loops by protecting cohesinSTAG1 from WAPL. Elife 9,.

Xue, M., Atallah, B. V. and Scanziani, M. (2014). Equalizing excitation-inhibition ratios across visual cortical neurons. Nature 511, 596–600.

Zakari, M., Yuen, K. and Gerton, J. L. (2015). Etiology and pathogenesis of the cohesinopathies. Wiley Interdiscip. Rev. Dev. Biol. 4, 489–504.

